# High-dimensional neuronal activity from low-dimensional latent dynamics: a solvable model

**DOI:** 10.1101/2025.06.03.657632

**Authors:** Valentin Schmutz, Ali Haydaroğlu, Shuqi Wang, Yixiao Feng, Matteo Carandini, Kenneth D. Harris

## Abstract

Computation in recurrent networks of neurons has been hypothesized to occur at the level of low-dimensional latent dynamics, both in artificial systems and in the brain. This hypothesis seems at odds with evidence from large-scale neuronal recordings in mice showing that neuronal population activity is high-dimensional. To demonstrate that low-dimensional latent dynamics and high-dimensional activity can be two sides of the same coin, we present an analytically solvable recurrent neural network (RNN) model whose dynamics can be exactly reduced to a low-dimensional dynamical system, but generates an activity manifold that has a high linear embedding dimension. This raises the question: Do low-dimensional latents explain the high-dimensional activity observed in mouse visual cortex? Spectral theory tells us that the covariance eigenspectrum alone does not allow us to recover the dimensionality of the latents, which can be low or high, when neurons are nonlinear. To address this indeterminacy, we develop Neural Cross-Encoder (NCE), an interpretable, nonlinear latent variable modeling method for neuronal recordings, and find that high-dimensional neuronal responses to drifting gratings and spontaneous activity in visual cortex can be reduced to low-dimensional latents, while the responses to natural images cannot. We conclude that the high-dimensional activity measured in certain conditions, such as in the absence of a stimulus, is explained by low-dimensional latents that are nonlinearly processed by individual neurons.

## 1 Introduction

The mammalian cortex comprises a large number of neurons, which, in principle, should allow it to use a high-dimensional neural code to represent sensory, motor, and cognitive information. Nevertheless, multi-neuronal recordings in nonhuman primates [1–4] have suggested that cortical populations perform computations by approximating low-dimensional dynamical systems [5, 6], with neuronal firing rates lying on a low-dimensional “neural manifold” [7]. In support of this hypothesis, low-dimensional dynamics have been inferred from multi-neuronal recordings through a wide variety of methods [8–23]; they spontaneously emerge in recurrent neural networks (RNNs) trained to solve behavioral tasks [2, 24–34]; and they appear in several theoretical models of noise-robust neuronal population dynamics [35–38]. A result that might at first sight challenge the low-dimensional dynamical systems hypothesis is that visual cortical population activity in mice has high linear dimension [39, 40] with shared neuronal covariance having a heavy-tailed eigenspectrum (see also [41] and [42] for recordings in cerebellum and across cortex, respectively). In particular, the shared covariance eigenspectrum has a power-law tail with an exponent close to 1 (*α* ≈ 1.04) [39] for responses to natural images and an exponent of *α* ≈ 1.14 for spontaneous activity [40]. Are these two views on the dimensionality of population activity compatible? Namely, can a low-dimensional dynamical system produce a neural manifold that has a high linear embedding dimension?

Here, we first construct a solvable RNN model that reconciles the low- and high-dimensional perspectives on population activity by carefully disambiguating the *linear* dimension of the system *before* and *after* the neurons’ nonlinearity, which we refer to as the pre- and post-activation dimension, respectively. This dichotomy refines the usual distinction between linear and “intrinsic” dimension [39, 43, 44], since the intrinsic dimension of a system is the same before and after any continuous, injective nonlinearity. Using the notions of pre- and post-activation linear dimensions, we show that our RNN can be exactly reduced to a low-dimensional dynamical system in the space of pre-activations, making the pre-activations low-dimensional. Then, we show that these latent dynamics generate high-dimensional post-activation activity that has a power-law covariance eigenspectrum. (In this work, dimension will always refer to linear dimension, unless stated otherwise.)

Before analyzing experimental recordings, we revisit the spectral theory of infinite-width neural networks (random feature kernels) [45–47] to quantitatively relate the pre-activation dimension, the neuronal activation function, and the post-activation covariance eigenspectrum. This three-way relationship tells us that high-dimensional activity is consistent with both low- and high-dimensional pre-activations. To uncover the pre-activation dimension of high-dimensional activity in visual cortex, we perform two-photon calcium recordings of tens of thousands of neurons from mouse visual cortex, and infer the pre-activation dimension using the Neural Cross-Encoder (NCE), an interpretable, nonlinear latent variable modeling method which models the activity of each neuron as a simple linear-nonlinear readout of low-dimensional latents. NCE reveals that both the responses to drifting gratings and spontaneous activity can be well approximated by low-dimensional pre-activations, but that responses to natural images cannot. This suggests that the encoding of natural images in visual cortex is already high-dimensional in the space of pre-activations.

## 2 Solvable RNN Model

To demonstrate how high-dimensional post-activations can arise from low-dimensional pre-activation dynamics, we first present a solvable RNN model whose autonomous dynamics is low-dimensional in the space of pre-activations, but high-dimensional in the space of post-activations, with the post-activations producing a power-law covariance eigenspectrum.

We consider an RNN consisting of *N* rate-units (neurons). The pre-activation *x*_*i*_ of neuron *i* evolves according to

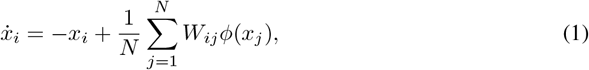

where *W*_*ij*_ denotes the synaptic weight from neuron *j* to neuron *i*, and *ϕ* : ℝ → ℝ _≥ 0_ is a nonlinear activation function converting the pre-activations into post-activations (firing rates). To define the weights *W*_*ij*_, we randomly place neurons on a ring [48–50] by assigning to each neuron *i* an independent and uniformly distributed angle *θ*_*i*_ ∈ [0, 2*π*) (Fig. 1A). The weights *W*_*ij*_ are then given by the following shifted cosine function:

**Figure 1.**
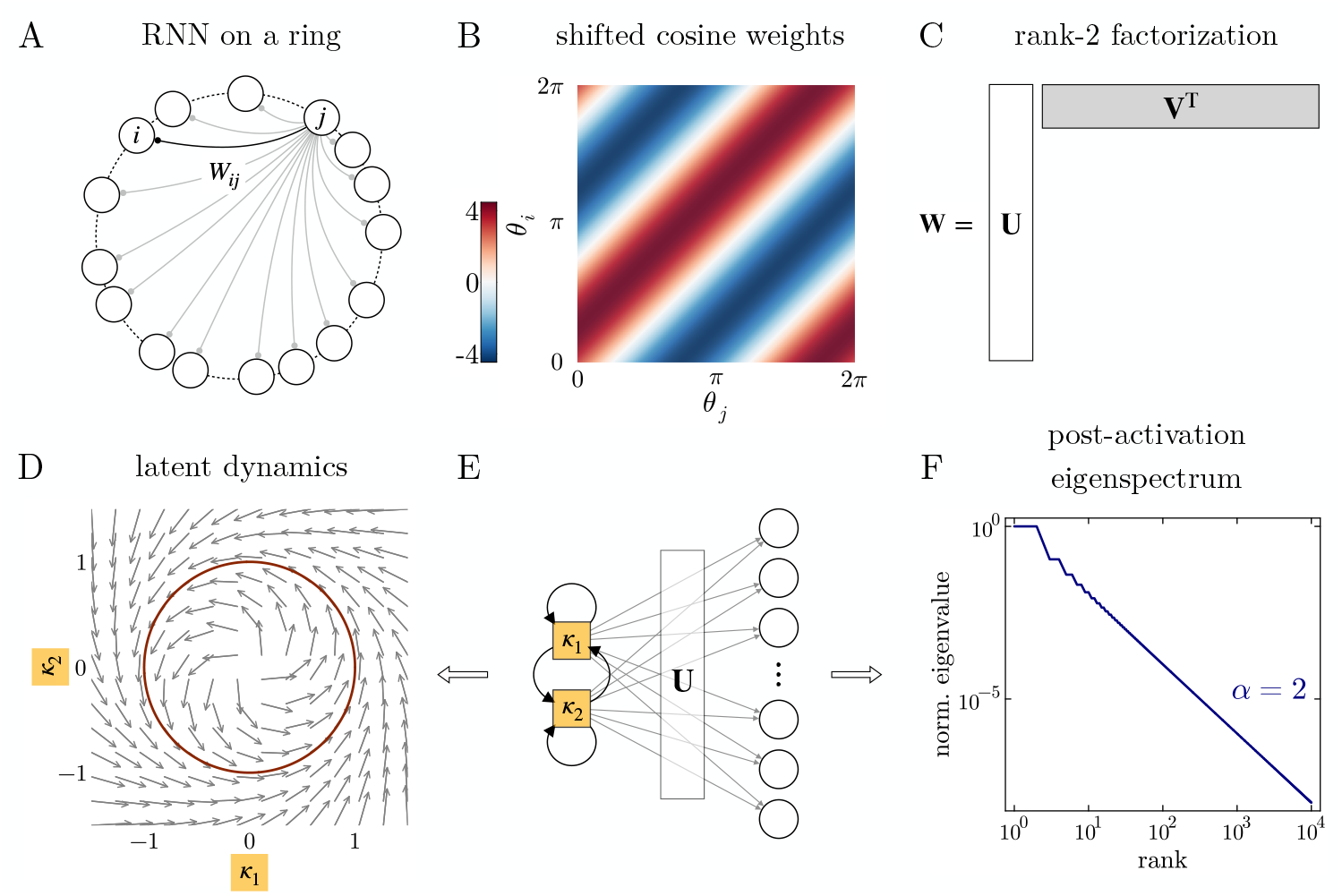
Solvable RNN model. (**A**) Schematic of the RNN model. (**B**) Shifted cosine function defining the position-dependent synaptic weights *W*_*ij*_. (**C**) Factorization of weight matrix **W** as the product of a 2-column matrix **U** and two-row matrix **V**^T^. (**D**) Vector field of the dynamics of the latent variables, *κ*_1_ and *κ*_2_, in the large-network limit. The latent dynamics produces a stable limit cycle on the unit circle. (**E**) Graphical representation of the RNN’s effective dynamics. (**F**) Post-activation covariance eigenspectrum. The eigenvalues follow a power-law with decay exponent *α* = 2. They are normalized such that the largest eigenvalue is 1. The activation function *ϕ* used is the step function defined in Eq. (3). Shown in D and F are theoretical values given by Eqs. (5) and (8), respectively, which closely match simulations of large networks (see Appendix C).

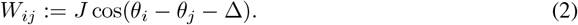

The shift Δ in Eq. (2) makes the weights asymmetric, with neurons sending their strongest excitatory output to neurons located at an angle Δ counter-clockwise (Fig. 1B). To make the model solvable, we assume that the activation function *ϕ* is the Heaviside step function Θ, i.e.,

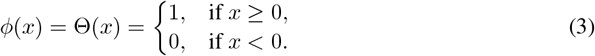

### 2.1 Low-dimensional pre-activation dynamics

The RNN model defined above has low-dimensional pre-activation dynamics because the weight matrix **W** has rank 2. Indeed, using an elementary trigonometric identity,^2^ **W** can be factorized as the outer product **W** = **UV**^T^ of two *N* × 2-matrices (Fig. 1C), with

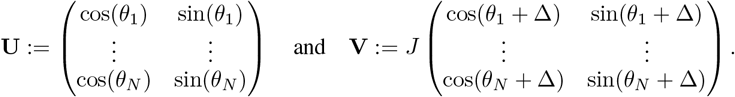

Then, following Beiran et al. [51], we can reduce the *N* -dimensional system, Eq. (1), to a 2-dimensional system describing the dynamics of the latent variables ***κ*** := **U**^†^**x**, where ^†^ denotes the pseudoinverse and **x** the *N* -dimensional vector of pre-activations (*x*_1_, …, *x*_*N*_)^T^. The dynamics of the latent variables follows

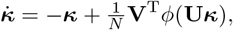

where *ϕ* is applied element-wise to the *N* -dimensional vector of pre-activations **U*κ*** = **x**. In the equation above, the vector *ϕ*(**U*κ***) = *ϕ*(**x**) represents the joint post-activations (firing rates) of the *N* neurons.

Taking the number of neurons *N* → ∞ yields a neural field limit [52] where the sum over neurons becomes an integral over the ring,

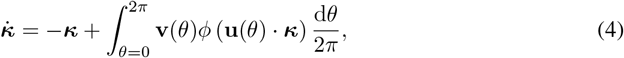

with **u**(*θ*) := (cos(*θ*), sin(*θ*))^T^ and **v**(*θ*) := *J*(cos(*θ* + Δ), sin(*θ* + Δ))^T^. Equation (4) describes the dynamics of the latent variables ***κ*** as the solution to a 2-dimensional dynamical system whose vector field involves an integral over the “circuit structure” [52] (the ring).

Since *ϕ* is the step function, the integral over the ring in Eq. (4) can be solved, and we obtain the solvable 2-dimensional dynamical system,

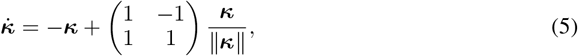

when 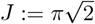 and Δ := *π/*4 (derivation presented in Appendix A). Equation (5) generates a stable limit cycle over the unit circle (Fig. 1D), which implies that the latent variables ***κ*** will eventually rotate on the unit circle indefinitely. In Appendix E, we provide other examples of low-rank RNNs for which the latent dynamics can be expressed in a tractable form similar to Eq. (5).

Neuronal activity, modeled here as the post-activations *ϕ*(**U*κ***), are simple *linear-nonlinear* readouts of latent variables ***κ*** (Fig. 1E). Hence, we have effectively reduced the dynamics of the RNN, Eq. (1), to a 2-dimensional latent dynamical system. We will say that neuronal activity is a *linear-nonlinear* function of latent variables if it is given by the composition of a linear mapping (**U**) and an element-wise, nondecreasing nonlinear mapping (*ϕ*).

### 2.2 Post-activations produce a power-law eigenspectrum

Since the dynamics of the latent variables is solvable in the large-network limit, and rotates on the unit circle, we can compute the correlation between the post-activations of two neurons. For any pair of neurons *i* and *j*, with positions *θ*_*i*_ and *θ*_*j*_ on the ring, respectively, the correlation of their post-activations *C*_*ij*_ is, in the long-recording limit, given by

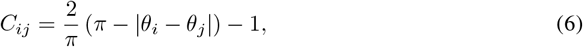

where |*θ*_*i*_− *θ*_*j*_| := cos^−1^(cos(*θ*_*i*_ − *θ*_*j*_)) is the absolute angle difference *θ*_*i*_ and *θ*_*j*_ (derivation presented in Appendix B).

We can find the eigenvalue spectrum of post-activations by noting that as the number of recorded neurons *M* → ∞, the eigenvalues of the *M* × *M* correlation matrix **C**^(*M*)^ defined by Eq. (6) converge to the eigenvalues of an integral operator. Since the angles *θ*_*i*_ are independently and uniformly sampled on circle [0, 2*π*), the correlation matrix **C**^(*M*)^ is a so-called Euclidean random matrix [53], that is, a matrix whose entries are given by the pairwise distances between randomly sampled points in a given space. Writing 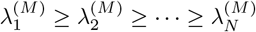 for the ranked eigenvalues of the matrix **C**^(*M*)^, random matrix theory [54, 55] tells us that, as *M* → ∞, the scaled eigenvalues 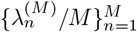, converge (in a ℓ_2_ sense) to the eigenvalues of the integral operator,

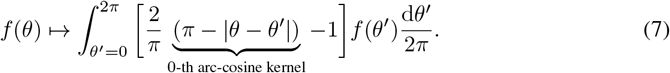

The eigenvalues of this integral operator can be computed analytically. In Eq. (7), we have highlighted the presence of the 0-th arc-cosine kernel *k*_0_(*θ, θ*^*′*^) := *π* − |*θ* − *θ*^*′*^| of Cho and Saul [45], which is well-known in machine learning, and whose eigenvalues have been computed in [46]. In short, by the rotational invariance of *k*_0_, we have that, for any positive integer *m*, the functions *θ* ↦ cos(*mθ*) and *θ* ↦ sin(*mθ*) are orthogonal eigenfunctions of the operator Eq. (7) sharing the same eigenvalue,

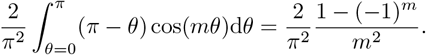

Using this result, we obtain that the ranked eigenvalues *λ*_1_ ≥ *λ*_2_ ≥ … of the operator Eq. (7) are given by

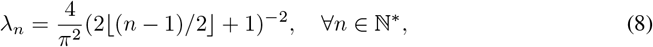

that is, eigenvalues come in identical pairs that decay exactly as a power law with decay exponent *α* = 2 (Fig. 1F). Hence, post-activations are high-dimensional in the sense that their covariance eigenspectrum has a heavy tail [39]. Although the model presented assumed, for simplicity, a cosine lateral connectivity on the ring, Eq. (2), similar results can be derived for more general lateral connectivity; see Appendix D for an example.

In summary, this solvable model shows that low-dimensional dynamics in the space of pre-activations can generate high-dimensional post-activations. The heavy tail of the covariance eigenspectrum implies that post-activations are not confined to any finite-dimensional linear subspace. Formally, the smallest vector space containing the post-activations generated by our model has the same size as the infinite-dimensional reproducing kernel Hilbert space associated with the kernel *k*_0_. We stress that, in this model, the heavy tail of the post-activation eigenspectrum is not due to noise, since we used a deterministic, non-chaotic RNN. Also, all the results presented above remain exact if the rate-units in Eq. (1) are replaced by linear-nonlinear-Poisson neurons, as spike noise cancels out in the limits we consider [38, 52].

## 3 Post-activation eigenspectrum depends on pre-activation dimension and activation function

To shed light on the relationship between the post-activation eigenspectrum, pre-activation dimension, and the activation function *ϕ*, we now turn to a more general setup, which allows us to relax some of the strong assumptions of the solvable model (Fig. 1E). First, we allow the number of latent variables *d* to be greater than 2, assuming that the latent variables, henceforth denoted by **z** (instead of ***κ***), are uniformly distributed on the unit sphere 𝕊^*d*−1^ in ℝ^*d*^. We assume that the pre-activations of the network are determined by passing the latent activity through a *N* × *d* feedforward weight matrix **U** with i.i.d. standard normal entries. In this setup (Fig. 2A), we call *d* the *pre-activation dimension*, as it sets the linear dimensionality of the pre-activations. In the solvable model of Sec. 2.2, for example, the pre-activation dimension was *d* = 2 (Fig. 1E). Finally, we replace the step function, Eq. (3), by the general rectified power activation function

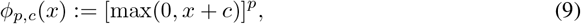

where the activation parameter *p* ∈ ℝ _≥0_ is a nonnegative real value and the bias *c* ∈ ℝ. (By convention, *ϕ*_0,*c*_(*x*) := Θ(*x* + *c*).)

**Figure 2.**
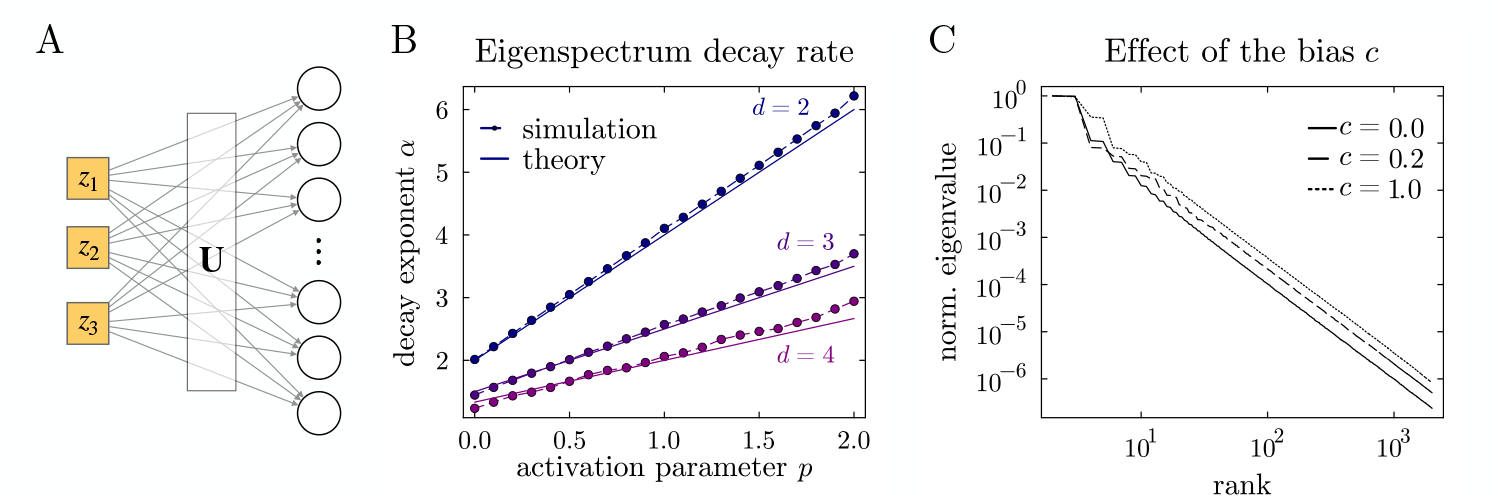
Eigenspectrum of the random feature kernel (11). (**A**) Schematic of the feedforward setup (in the case of three input variables). (**B**) Comparison between simulations of the eigenspectrum decay rate *α* and the theoretical value predicated by Conjecture 1, for *p* ∈ [0, 2] and *d* = 2, 3, 4. Up to finite-size effects (number of neurons *N* = 2 *·* 10^4^ and number of inputs *T* = 10^4^), simulations match the theoretical prediction 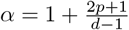. (In this plot, *c* = 0.) (**C**) Simulations of the eigenspectrum for various bias parameter *c*, while keeping *p* and *d* fixed. (In this plot, *p* = 0 and *d* = 2, which gives *α* = 2. Eigenvalues are normalized as in Fig. 1F.)

This setup can be analyzed within the framework of random feature kernels (see [56, Sec. 9.5]). Denoting *µ*_*d*−1_ the uniform probability measure on the sphere 𝕊^*d*−1^, let us take *T* independent latent variable samples **z**_1_, …, **z**_*T*_ from *µ*_*d*− 1_, and define the *N* × *T* post-activation matrix **A**^(*N,T*)^ := (*ϕ*_*p,c*_(**Uz**_1_), …, *ϕ*_*p,c*_(**Uz**_*T*_)). In the limits *N* → ∞ and *T* → ∞ taken successively, the covariance eigenspectrum of **A**^(*N,T*)^ converges (when properly scaled) to the eigenvalue spectrum of the integral operator

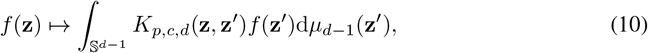

where *K*_*p,c,d*_ : 𝕊^*d*−1^ × 𝕊^*d*−1^ → ℝ is the *random feature kernel*

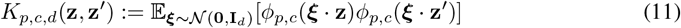

(see [46, 56] or Appendix F for more details).

Drawing intuition from Fourier analysis, the smoothness of a function (here the kernel) should be related to the decay rate of its Fourier transform (here the eigenspectrum)—the smoother the function, the faster the decay rate of its Fourier transform. Known results on the eigenvalues of random feature kernels for the cases *p* = 0 and *p* = 1, with *c* = 0, confirm this intuition and show how it extends to general integers *d* [46]. Extrapolating those results to any nonnegative *p* and any real *c*, we get the following conjecture.

### Conjecture 1.

*For any p* ∈ ℝ _≥0_, *c* ∈ ℝ, *and any integer d* ≥ 2, *the ranked eigenvalues λ*_1_ ≥ *λ*_2_ ≥ … *of the integral operator* (10) *obey the following power-law decay:*

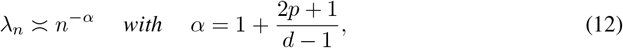

*where a*_*n*_ ≍ *b*_*n*_ *means* lim_*n*→+∞_ *a*_*n*_*/b*_*n*_ = *C* ∈ (0, +∞).

To the best of our knowledge, Conjecture 1 is not a straightforward consequence of any existing result in theoretical machine learning [47, 57, 58] or harmonic analysis [59–62], hence our presentation of Eq. (12) as a conjecture. Note that when the activation parameter *p* is an integer, *ϕ*_*p,c*_ is *p*-times weakly differentiable, that is, the first *p* weak derivatives^3^ of *ϕ*_*p,c*_ are all locally integrable. This, and the fact that the bias *c* does not affect the decay rate, suggest a further extension of the conjecture to more general activation functions, with *p* replaced by the weak differentiability of the activation function.

We tested Conjecture 1 numerically by performing PCA on large post-activation matrices **A**^(*N,T*)^. The linear and continuous dependence of the decay exponent *α* on the activation parameter *p* predicted by Eq. (12) was confirmed in simulations (Fig. 2B). Simulations also confirmed that the bias *c* of the activation function does not affect the decay rate (Fig. 2C), a fact already mentioned in [63, 64].

To summarize, the spectral theory of random feature kernels suggests a three-way relationship between the power-law tail exponent of the post-activation eigenspectrum, the pre-activation dimension, and the activation function. This relationship should hold when we can consider the neurons as linear-nonlinear functions of the latent vector, with weights that vary randomly and independently between neurons. The relationship suggests that, when high-dimensional neuronal activity (modeled here as post-activations) is observed [39, 40], two scenarios are possible: high-dimensional activity could arise from nonlinear transformation of low-dimensional latent states, or it could reflect pre-activations that are already high-dimensional. To distinguish these two scenarios, we propose, in what follows, a method for inferring the pre-activation dimension of neuronal activity in experimental recordings.

## 4 Latent Variable Modeling of Neuronal Recordings

To estimate the input dimensionality of neuronal activity in mouse visual cortex (as defined in Sec. 3) we developed the Neural Cross-Encoder (NCE), a nonlinear generalization of Reduced Rank Regression. Using NCE, we show that high-dimensional neuronal responses to drifting gratings are well-approximated by a linear-nonlinear readout of a low dimensional latent variable, whereas responses to natural images are not. Finally, we apply NCE to high-dimensional spontaneous dynamics in the cortex and find that they are well-approximated by a linear-nonlinear readout of low-dimensional latents.

### 4.1 Experimental Data

We conducted large-scale volumetric two-photon microscopy on awake, adult mice during visual stimulation and spontaneous activity. We targeted primary and higher visual cortices with a Light Beads Microscope [65], and extracted deconvolved activity traces for 19,223 ± 2,948 neurons using Suite3D [66] as described in Appendix H. Recordings were performed in three stimulus conditions: (1) responses to 320 full-field drifting grating stimuli with 2-14 repeats each; (2) responses to 1866 natural images with 2 repeats each; (3) spontaneous activity in the absence of stimuli for 10-15 minutes.

### 4.2 Neural Cross-Encoder (NCE)

The Neural Cross-Encoder (NCE) divides neurons randomly into two sets: a source set and a target set. It predicts the activity **b**_*t*_ of the target set from the source set **a**_*t*_ via a non-linear readout of a set of latents, **z**_*t*_ (Fig. 3A). NCE uses a multi-layer feedforward encoder ℰ that ends in a bottleneck layer whose activity **z**_*t*_ = ℰ (**a**_*t*_) represents a low-dimensional latent state estimated from the source neurons. The reason to use this rather than an autoencoder, which predicts one set of neurons from themselves, is to discard variability that is not shared across neurons. The NCE we used here has a single power-ReLU output layer, matching the setup of section 3, and a 3-layer encoder allowing flexible estimation of latent variables, so that the number of latent variables can be readily interpreted as the pre-activation dimension. We train NCE with stochastic gradient descent on source-target activity pairs as described in Appendix I. When all nonlinearities are removed, NCE becomes equivalent to Reduced Rank Regression [67]. When predicting stimulus-driven activity, we pair the activity of source and target neurons on different repeats of the same stimulus, to also discard shared variability that is not related to the stimulus [39].

**Figure 3.**
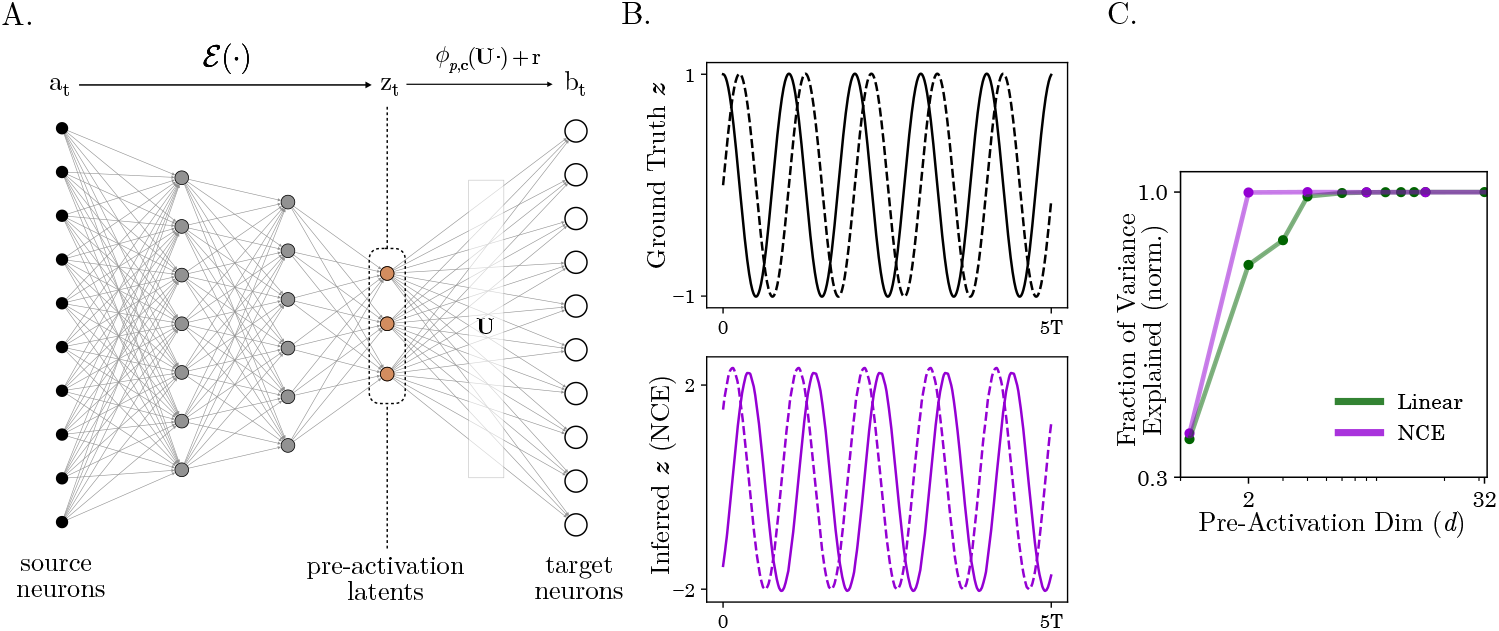
Validation of the Neural Cross Encoder. (**A**) Schematic of the NCE. **B** Simulated pre-activation latents of a two-dimensional toy model (top), and the inferred pre-activation latents using NCE (bottom). **C** Reconstruction score as a function of pre-activation dimensionality for Linear (RRR) and NCE models on simulated data.

#### Linear-nonlinear readout

The recorded activity of a set of target neurons at time 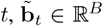, is modeled as a weighted sum of the latent variables, **z**_*t*_ ∈ ℝ^*d*^, passed through a nondecreasing nonlinearity:

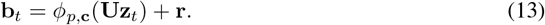

Here, *ϕ*_*p*,**c**_ is the rectified power activation function, defined in Eq. 9, with a power parameter *p* that is constant across neurons, a pre-activation bias that varies across neurons encoded by an *N* -dimensional vector **c**, and a post-activation added bias **r** to account for non-zero baseline firing rates. The decoder parameters {*p*, **c, U, r**} are learned alongside the encoder parameters of ℰ. The fact that the decoder, Eq. (13), has a single-layer is crucial as it allows us to interpret the latents (**z**_*t*_) as linear factors of the observed neurons’ pre-activations (**Uz**_*t*_). It is this constrained decoder that allows us to infer the pre-activation dimension of neuronal activity; in comparison, a multi-layer decoder as used in [13, 23] would infer something closer to the intrinsic dimension of neuronal activity, which is not our goal.

### 4.3 Results

#### NCE identifies the latent dimensionality of simulated data

To validate that NCE can identify the pre-activation latent variables, we test it on simulated data generated from the toy model in Sec. 2 with a ReLU readout (*p* = 1) and *d* = 2. NCE recovers the true latents up to a scaling and a shift (Fig 3B). Moreover, NCE can explain all of the variance in the population with only two pre-activation dimensions, while the corresponding linear model (Reduced Rank Regression) requires more dimensions (Fig 3C).

#### Pre-activation dimension is low for grating responses, high for natural image responses

We next consider the pre-activation dimensional of visual stimulus responses of visual cortex neurons. To ensure that the NCE focused on the stimulus responses, and not correlated ongoing activity such as spontaneous activity or encoding of movements, the activity of the target cells and the source cells were taken from different repeats of the same stimuli. In the case of drifting gratings, for which we know there is a low-dimensional latent variable (the grating orientation), an NCE model with low-dimensional pre-activations accurately predicts neuronal responses (Fig. 4A,B). NCE requires fewer dimensions (5.5 ± 1.2, mean ± std) than the corresponding linear model (13.9 ± 3.0) to predict 95% of the explainable variance (defined as the maximum variance explained across all *d* in both models). On the other hand, NCE models with low pre-activation dimension are not sufficient to predict responses to natural images (Fig. 4C,D), requiring 93.9 ± 6.0 dimensions to reach the threshold, suggesting that natural images produce high-dimensional representations in the space of pre-activations (but see Limitations below). Linear models only account for a smaller fraction of the total variance (Fig. 4D), and therefore underestimate the dimensionality of natural image responses (48.0 ± 13.1).

**Figure 4.**
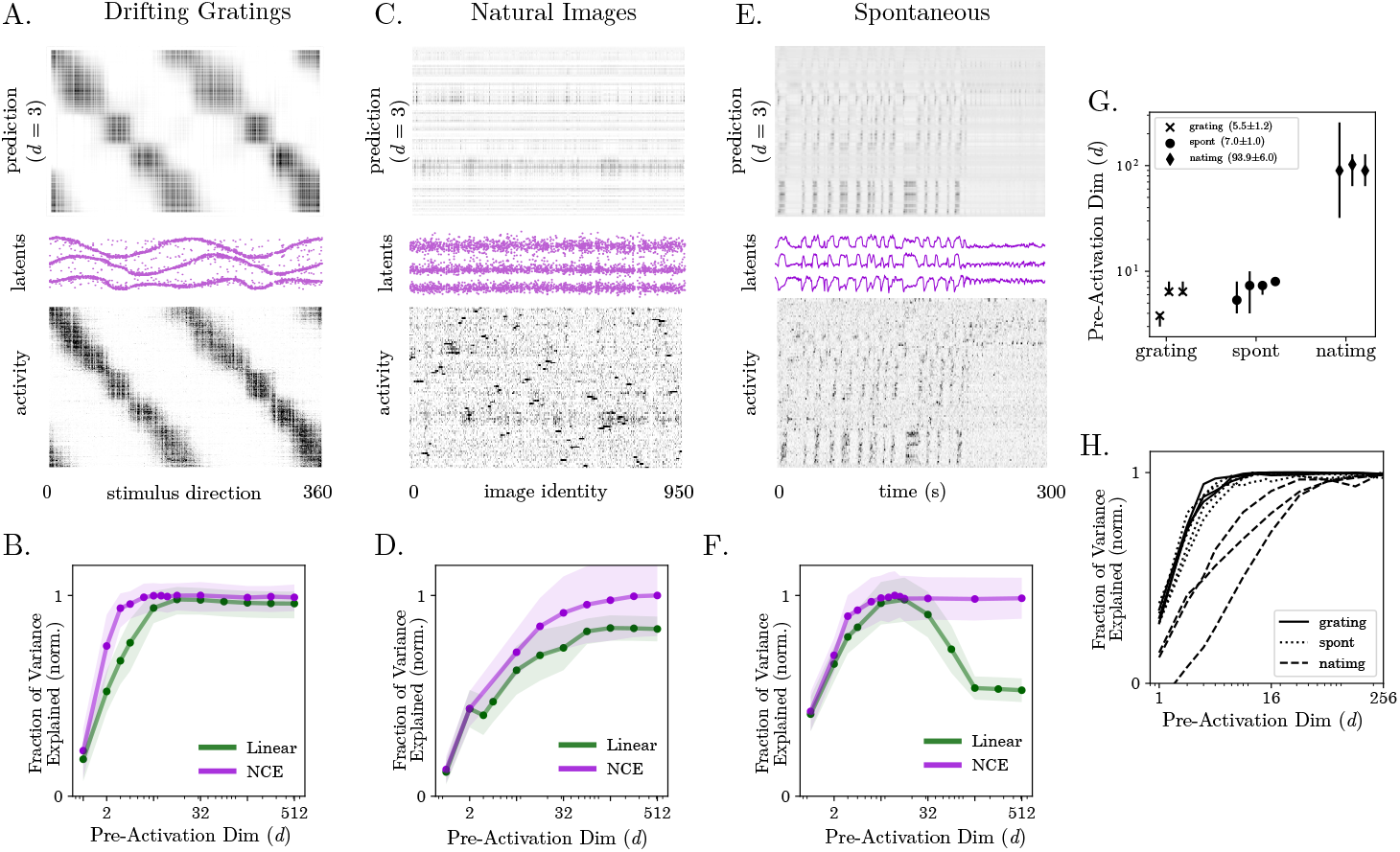
Pre-activation dimension in visual cortex depends on stimulus condition. (**A**) Rasters of true (bottom) and predicted (top) neuronal activity, and estimated latents (middle) for 1000 example neurons using an NCE with pre-activation dimension *d* = 3, for drifting grating responses. (**B**) Fraction of variance explained as a function of pre-activation dimension for an example session of grating responses. Plotted values are the mean across source/target selections for NCE (purple) and Reduced Rank Regression (green). Shaded regions are the standard deviation across source/target selections. All experiments contain 500 source and 1000 target neurons, and all scores are computed on a held-out test set. (**C**,**D**) Same as **A**,**B** for natural image responses. (**E**,**F**) Same as **A**,**B** for spontaneous activity. (**G**) Estimated pre-activation dimension *d* sufficient to capture 95% of explainable variance in the NCE across the three conditions. Each point is the mean for a single imaging session, error bars indicate the min/max across source/target selections. (**H**) Fraction of variance explained by NCE models with varying pre-activation dimension (log scale) across the three conditions. Each line represents a single imaging session.

#### Spontaneous activity has low pre-activation dimension

Spontaneous activity is well predicted by NCE with low pre-activation dimension (Fig. 4E,F). Across all recordings, spontaneous activity of 1000 target neurons has an estimated pre-activation dimension *d* of 7.0 ± 1.0 (mean ± std.), somewhat larger than grating responses but substantially lower than natural images responses (Fig. 4G,H). The linear model finds a similar dimension (7.5 ± 2.0), though its performance deteriorates at high pre-activation dimensions due to overfitting, while the NCE performance remains consistent (Fig. 4F).

These results indicate that visual cortex activity can be modeled as a linear-nonlinear transformation of a latent vector, which is low-dimensional for grating responses and spontaneous activity, but high-dimensional for natural image responses. In the case of grating responses, we find latents that resemble the sine and cosine of the stimulus angle (Fig. 4A)—this is what one would expect to find if neurons follow the canonical model of simple cells in visual cortex [68]. On the other hand, during spontaneous activity, the dynamics of the latent variables are correlated with the running speed of the mouse, and are perhaps related to its arousal state (see Appendix I.4).

## 5 Summary of Technical Contributions and Previous Works

### Latent dynamics of low-rank RNNs

Low-rank RNNs are tractable models of how the brain can perform computations through low-dimensional population dynamics [36, 51, 69–72]. In particular, the dynamics of certain low-rank RNNs, in the large-network limit, reduce to that of “effective circuits”, i.e., dynamical systems describing the evolution of the latent variables [51, 52, 73]. A limitation of these effective dynamical systems is that the expression of the vector field involves an integral over the distribution of weights (the “circuit structure” [52]), making them somewhat opaque and costly to solve numerically in general. In this work (Sec. 2.1 and Appendix E), we prove that, in several special cases that are beyond the case treated in [74], the integral over the weight distribution can be solved, yielding simple exact equations for the latent dynamics.

### Eigenvalue decay of random feature kernels

In the infinite-width limit, two-layer neural networks with random input weights behave like *random feature kernels* that depend on the distribution of input weights and the activation function of the neurons in the hidden layer [56, Sec. 9.5]. This functional perspective can be generalized to deep networks [45, 75] and constitute the basis of the Neural Tangent Kernel formalism for studying learning dynamics [76]. When the activation function is the ReLU, the decay rate of the eigenvalues has been proven to be polynomial [46, 47, 58, 64, 77, 78], even when inputs are not assumed to be uniformly distributed on the sphere [58]; for general results on dot-product kernels, see [57, 61, 63, 79, 80]. In this work, we propose a simple formula that links the power-law exponent of the eigenvalue decay rate, the power of the rectified-power activation function, and the input dimensionality. This formula, which goes beyond known results [46, 47, 64], is presented as a conjecture that we test in simulations.

### Latent variable modeling of neuronal activity

While most latent variable models of neuronal activity were originally developed for electrophysiological recordings [8–23], some are tailored for calcium recordings [81–83]. These models vary in their mechanistic interpretability: The inferred latents are either abstract variables, for example when the model’s mapping from latents to neuronal activity involves a multi-layer neural network [13, 23], or they can be interpreted as linear factors of the neurons’ pre-activations, as in [9, 10]. With nonlinear dimensionality reduction methods such as CEBRA [84] or Rastermap [85], latent variables are also abstract as there is no explicit mapping going from the latents to neuronal activity. In this work (Sec. 4), we developed NCE, a latent variable model for calcium recordings that models neuronal activity as an interpretable linear-nonlinear readout of latent variables. NCE also uses a cross-encoding scheme, which allows it to discard variability not shared across neurons. We demonstrate that NCE is capable of identifying a low-dimensional pre-activation space even when the recorded neuronal activity has high linear dimension.

## 6 Discussion

### Dimensionality of neural systems

The solvable RNN model we proposed produces cyclic population dynamics that is low-dimensional in the space of pre-activations, and high-dimensional in the space of post-activations (firing rates). Thus, an RNN can produce trajectories that are simultaneously low-dimensional and high-dimensional, depending on the variables being considered. In this work, we focused on the notion of *linear* dimensionality and adopted an infinite-dimensional Hilbert space formalism borrowed from kernel methods to characterize the linear dimensionality of firing rate trajectories in the large-network limit. Of course, the “intrinsic” dimension of neuronal activity is equal to 1, since the dynamics is periodic; this highlights the important distinction between intrinsic and linear (or “embedding”) dimension of neuronal activity (see also [43, 44]). The type of high-dimensional activity our model produces is computationally relevant: It can be exploited by a downstream readout neuron to represent arbitrary periodic functions (see also [86]), or, following the random readout approach of [87, 88], it can be used to represent a Gaussian process prior over periodic functions (see Appendix G).

Two definitions of high-dimensional neuronal activity have been studied in neuroscience. Throughout this work, we define high-dimensional neuronal activity to mean a covariance eigenspectrum with a heavy tail that decays strictly faster that 1*/n* [39]. This definition is well-suited for systems generating activity whose pairwise correlations do not converge to zero in the limit of large network size. In contrast, random chaotic RNNs––solvable models whose pairwise correlations do converge to zero––produce a different form of high-dimensional activity, where the eigenspectrum decays slower than 1*/n* [89], reflecting noise that is not shared between neurons. While a comprehensive comparison of these two types of high-dimensional activity is beyond the scope of this work, we mention that the latter is relevant when one wants to study the noisiness of neuronal responses [90].

### Pre-activations and subthreshold membrane potentials

We developed NCE to disentangle the linear dimension of neuronal activity and of the neurons’ pre-activations. We show that even when activity is high-dimensional, it can be well-explained with low-dimensional pre-activations in the case of grating responses and spontaneous activity. From a biological point of view, how should we interpret the pre-activations inferred by NCE? If one assumes that the link between synaptic integration and neuronal firing is well approximated by a simple nonlinear activation function in cortical neurons, as NCE does, one could argue that pre-activations represent estimates of the neurons’ synaptic inputs or subthreshold membrane potentials. This is an experimental prediction that large-scale voltage imaging of neuronal populations [91] may make testable in the near future.

### Limitations

Training NCE, which is a non-convex optimization problem, can be challenging when neural data is limited. The duration of imaging experiments is limited to 3.5 h, which yields only a few thousand training examples per session for tens of thousands of neurons; thus, optimization can get stuck in local minima. To facilitate training, we limit to fitting only 1000 highly responsive neurons at a time, and use tools such as data augmentation and pretraining as described in Appendix I. The fact that we were not able to accurately predict neuronal responses to natural images with a low-dimensional NCE model does not necessarily exclude the possibility for such a model to exist, and one could possibly find it with a larger training set or a different model class/hyperparameter configuration. Note that this work did not revisit the problem of how to estimate the tail of the shared covariance eigenspectrum from neuronal recordings, which is discussed in [39, 40, 92, 93].

## Acknowledgments and Disclosure of Funding

The authors thank Louis Pezon for useful discussions; Michael Krumin, Bex Terry and Charu B. Reddy for experimental support; Kimberly Ren for feedback on the manuscript. We also thank the four anonymous reviewers who have helped improve the manuscript. This work was funded by UKRI (Frontier Award EP/X022366/1 to MC), BBSRC (grant BB/W019884/1 to MC), the National Institutes of Health BRAIN initiative (grant U01NS126057 to MC), the Wellcome Trust (Investigator Award 223144/Z/21/Z to MC and KDH), and the ERC (101097874 to KDH). MC holds the GlaxoSmithKline / Fight for Sight Chair in Visual Neuroscience. AH is supported by a studentship from the Gatsby Charitable Foundation (GAT3755) and the Wellcome Trust (219627/Z/19/Z). VS is supported by a Royal Society Newton International Fellowship (NIF\R1\231927) and a fellowship from the Swiss National Science Foundation (grant no. 222150).

## NeurIPS Paper Checklist

### 1. Claims

Question: Do the main claims made in the abstract and introduction accurately reflect the paper’s contributions and scope?

Answer: [Yes]

Justification: The claims we make in abstract and introduction result our results. Guidelines:

- The answer NA means that the abstract and introduction do not include the claims made in the paper.
- The abstract and/or introduction should clearly state the claims made, including the contributions made in the paper and important assumptions and limitations. A No or NA answer to this question will not be perceived well by the reviewers.
- The claims made should match theoretical and experimental results, and reflect how much the results can be expected to generalize to other settings.
- It is fine to include aspirational goals as motivation as long as it is clear that these goals are not attained by the paper.

### 2. Limitations

Question: Does the paper discuss the limitations of the work performed by the authors? Answer: [Yes]

Justification: There is a paragraph titled “Limitations” in the discussion section where we clearly state the limitations of our results.

Guidelines:

- The answer NA means that the paper has no limitation while the answer No means that the paper has limitations, but those are not discussed in the paper.
- The authors are encouraged to create a separate “Limitations” section in their paper.
- The paper should point out any strong assumptions and how robust the results are to violations of these assumptions (e.g., independence assumptions, noiseless settings, model well-specification, asymptotic approximations only holding locally). The authors should reflect on how these assumptions might be violated in practice and what the implications would be.
- The authors should reflect on the scope of the claims made, e.g., if the approach was only tested on a few datasets or with a few runs. In general, empirical results often depend on implicit assumptions, which should be articulated.
- The authors should reflect on the factors that influence the performance of the approach. For example, a facial recognition algorithm may perform poorly when image resolution is low or images are taken in low lighting. Or a speech-to-text system might not be used reliably to provide closed captions for online lectures because it fails to handle technical jargon.
- The authors should discuss the computational efficiency of the proposed algorithms and how they scale with dataset size.
- If applicable, the authors should discuss possible limitations of their approach to address problems of privacy and fairness.
- While the authors might fear that complete honesty about limitations might be used by reviewers as grounds for rejection, a worse outcome might be that reviewers discover limitations that aren’t acknowledged in the paper. The authors should use their best judgment and recognize that individual actions in favor of transparency play an important role in developing norms that preserve the integrity of the community. Reviewers will be specifically instructed to not penalize honesty concerning limitations.

### 3. Theory assumptions and proofs

Question: For each theoretical result, does the paper provide the full set of assumptions and a complete (and correct) proof?

Answer: [Yes]

Justification: Our nontrivial mathematical statements are either fully proved or clearly stated as conjectures (e.g. Conjecture 1 in the main text).

Guidelines:

- The answer NA means that the paper does not include theoretical results.
- All the theorems, formulas, and proofs in the paper should be numbered and cross-referenced.
- All assumptions should be clearly stated or referenced in the statement of any theorems.
- The proofs can either appear in the main paper or the supplemental material, but if they appear in the supplemental material, the authors are encouraged to provide a short proof sketch to provide intuition.
- Inversely, any informal proof provided in the core of the paper should be complemented by formal proofs provided in appendix or supplemental material.
- Theorems and Lemmas that the proof relies upon should be properly referenced.

### 4. Experimental result reproducibility

Question: Does the paper fully disclose all the information needed to reproduce the main experimental results of the paper to the extent that it affects the main claims and/or conclusions of the paper (regardless of whether the code and data are provided or not)?

Answer: [Yes]

Justification: The architecture, training procedure and hyperparameters for the Neural Cross-Encoder are described in detail. While the text descriptions are sufficient to reproduce the results, we also provide the code for *in silica* experiments. We also share the preprocessed neuronal datasets. The methods for *in vivo* experiments and the preprocessing of imaging data are described clearly to enable reproducibility.

Guidelines:

- The answer NA means that the paper does not include experiments.
- If the paper includes experiments, a No answer to this question will not be perceived well by the reviewers: Making the paper reproducible is important, regardless of whether the code and data are provided or not.
- If the contribution is a dataset and/or model, the authors should describe the steps taken to make their results reproducible or verifiable.
- Depending on the contribution, reproducibility can be accomplished in various ways. For example, if the contribution is a novel architecture, describing the architecture fully might suffice, or if the contribution is a specific model and empirical evaluation, it may be necessary to either make it possible for others to replicate the model with the same dataset, or provide access to the model. In general. releasing code and data is often one good way to accomplish this, but reproducibility can also be provided via detailed instructions for how to replicate the results, access to a hosted model (e.g., in the case of a large language model), releasing of a model checkpoint, or other means that are appropriate to the research performed.
- While NeurIPS does not require releasing code, the conference does require all submissions to provide some reasonable avenue for reproducibility, which may depend on the nature of the contribution. For example
  a. If the contribution is primarily a new algorithm, the paper should make it clear how to reproduce that algorithm.
  b. If the contribution is primarily a new model architecture, the paper should describe the architecture clearly and fully.
  c. If the contribution is a new model (e.g., a large language model), then there should either be a way to access this model for reproducing the results or a way to reproduce the model (e.g., with an open-source dataset or instructions for how to construct the dataset).
  d. We recognize that reproducibility may be tricky in some cases, in which case authors are welcome to describe the particular way they provide for reproducibility. In the case of closed-source models, it may be that access to the model is limited in some way (e.g., to registered users), but it should be possible for other researchers to have some path to reproducing or verifying the results.

### 5. Open access to data and code

Question: Does the paper provide open access to the data and code, with sufficient instructions to faithfully reproduce the main experimental results, as described in supplemental material?

Answer: [Yes]

Justification: All code required to reproduce the main experimental results is anonymized and submitted with the supplementary materials. Preprocessed neural datasets, comprised of deconvolved spontaneous activity and averaged, deconvolved stimulus responses, is also anonymized and submitted with the supplementary materials.

Guidelines:

- The answer NA means that paper does not include experiments requiring code.
- Please see the NeurIPS code and data submission guidelines (https://nips.cc/public/guides/CodeSubmissionPolicy) for more details.
- While we encourage the release of code and data, we understand that this might not be possible, so “No” is an acceptable answer. Papers cannot be rejected simply for not including code, unless this is central to the contribution (e.g., for a new open-source benchmark).
- The instructions should contain the exact command and environment needed to run to reproduce the results. See the NeurIPS code and data submission guidelines (https://nips.cc/public/guides/CodeSubmissionPolicy) for more details.
- The authors should provide instructions on data access and preparation, including how to access the raw data, preprocessed data, intermediate data, and generated data, etc.
- The authors should provide scripts to reproduce all experimental results for the new proposed method and baselines. If only a subset of experiments are reproducible, they should state which ones are omitted from the script and why.
- At submission time, to preserve anonymity, the authors should release anonymized versions (if applicable).
- Providing as much information as possible in supplemental material (appended to the paper) is recommended, but including URLs to data and code is permitted.

### 6. Experimental setting/details

Question: Does the paper specify all the training and test details (e.g., data splits, parameters, how they were chosen, type of optimizer, etc.) necessary to understand the results?

Answer: [Yes]

Justification: All relevant details of the training and testing of presented models are described in the main text and appendices, including the dataset generation, hyperparameters, and optimization procedure.

Guidelines:

- The answer NA means that the paper does not include experiments.
- The experimental setting should be presented in the core of the paper to a level of detail that is necessary to appreciate the results and make sense of them.
- The full details can be provided either with the code, in appendix, or as supplemental material.

### 7. Experiment statistical significance

Question: Does the paper report error bars suitably and correctly defined or other appropriate information about the statistical significance of the experiments?

Answer: [Yes]

Justification: On Fig. 4, error bars are reported and clearly defined. The factors of variability they represent, including variability from subselection of neurons within a session and variability across session, are clearly stated. On Fig. 2B, error bars are not reported as the number of points representing single simulations (*>* 20 per line) already visually indicates the variance of the simulations.

Guidelines:

- The answer NA means that the paper does not include experiments.
- The authors should answer “Yes” if the results are accompanied by error bars, confidence intervals, or statistical significance tests, at least for the experiments that support the main claims of the paper.
- The factors of variability that the error bars are capturing should be clearly stated (for example, train/test split, initialization, random drawing of some parameter, or overall run with given experimental conditions).
- The method for calculating the error bars should be explained (closed form formula, call to a library function, bootstrap, etc.)
- The assumptions made should be given (e.g., Normally distributed errors).
- It should be clear whether the error bar is the standard deviation or the standard error of the mean.
- It is OK to report 1-sigma error bars, but one should state it. The authors should preferably report a 2-sigma error bar than state that they have a 96% CI, if the hypothesis of Normality of errors is not verified.
- For asymmetric distributions, the authors should be careful not to show in tables or figures symmetric error bars that would yield results that are out of range (e.g. negative error rates).
- If error bars are reported in tables or plots, The authors should explain in the text how they were calculated and reference the corresponding figures or tables in the text.

### 8. Experiments compute resources

Question: For each experiment, does the paper provide sufficient information on the computer resources (type of compute workers, memory, time of execution) needed to reproduce the experiments?

Answer: [Yes]

Justification: All experiments are undertaken on a single workstation, with a total compute time no greater than 7 days. Further details of the workstation and compute time are provided in the appendices

Guidelines:

- The answer NA means that the paper does not include experiments.
- The paper should indicate the type of compute workers CPU or GPU, internal cluster, or cloud provider, including relevant memory and storage.
- The paper should provide the amount of compute required for each of the individual experimental runs as well as estimate the total compute.
- The paper should disclose whether the full research project required more compute than the experiments reported in the paper (e.g., preliminary or failed experiments that didn’t make it into the paper).

### 9. Code of ethics

Question: Does the research conducted in the paper conform, in every respect, with the NeurIPS Code of Ethics https://neurips.cc/public/EthicsGuidelines?

Answer: [Yes]

Justification: The research presented in the paper conforms with the NeurIPS Code of Ethics. Guidelines:

- The answer NA means that the authors have not reviewed the NeurIPS Code of Ethics.
- If the authors answer No, they should explain the special circumstances that require a deviation from the Code of Ethics.
- The authors should make sure to preserve anonymity (e.g., if there is a special consideration due to laws or regulations in their jurisdiction).

### 10. Broader impacts

Question: Does the paper discuss both potential positive societal impacts and negative societal impacts of the work performed?

Answer: [NA]

Justification: This is a foundational research paper in the field of computational and systems neuroscience.

Guidelines:

- The answer NA means that there is no societal impact of the work performed.
- If the authors answer NA or No, they should explain why their work has no societal impact or why the paper does not address societal impact.
- Examples of negative societal impacts include potential malicious or unintended uses (e.g., disinformation, generating fake profiles, surveillance), fairness considerations (e.g., deployment of technologies that could make decisions that unfairly impact specific groups), privacy considerations, and security considerations.
- The conference expects that many papers will be foundational research and not tied to particular applications, let alone deployments. However, if there is a direct path to any negative applications, the authors should point it out. For example, it is legitimate to point out that an improvement in the quality of generative models could be used to generate deepfakes for disinformation. On the other hand, it is not needed to point out that a generic algorithm for optimizing neural networks could enable people to train models that generate Deepfakes faster.
- The authors should consider possible harms that could arise when the technology is being used as intended and functioning correctly, harms that could arise when the technology is being used as intended but gives incorrect results, and harms following from (intentional or unintentional) misuse of the technology.
- If there are negative societal impacts, the authors could also discuss possible mitigation strategies (e.g., gated release of models, providing defenses in addition to attacks, mechanisms for monitoring misuse, mechanisms to monitor how a system learns from feedback over time, improving the efficiency and accessibility of ML).

### 11. Safeguards

Question: Does the paper describe safeguards that have been put in place for responsible release of data or models that have a high risk for misuse (e.g., pretrained language models, image generators, or scraped datasets)?

Answer: [Yes]

Justification: Our results and data analysis method have no risk of being misused. Guidelines:

- The answer NA means that the paper poses no such risks.
- Released models that have a high risk for misuse or dual-use should be released with necessary safeguards to allow for controlled use of the model, for example by requiring that users adhere to usage guidelines or restrictions to access the model or implementing safety filters.
- Datasets that have been scraped from the Internet could pose safety risks. The authors should describe how they avoided releasing unsafe images.
- We recognize that providing effective safeguards is challenging, and many papers do not require this, but we encourage authors to take this into account and make a best faith effort.

### 12. Licenses for existing assets

Question: Are the creators or original owners of assets (e.g., code, data, models), used in the paper, properly credited and are the license and terms of use explicitly mentioned and properly respected?

Answer: [Yes]

Justification: Open-source software packages used in the work used within the terms of their licenses and credited in the Appendix.

Guidelines:

- The answer NA means that the paper does not use existing assets.
- The authors should cite the original paper that produced the code package or dataset.
- The authors should state which version of the asset is used and, if possible, include a URL.
- The name of the license (e.g., CC-BY 4.0) should be included for each asset.
- For scraped data from a particular source (e.g., website), the copyright and terms of service of that source should be provided.
- If assets are released, the license, copyright information, and terms of use in the package should be provided. For popular datasets, paperswithcode.com/datasets has curated licenses for some datasets. Their licensing guide can help determine the license of a dataset.
- For existing datasets that are re-packaged, both the original license and the license of the derived asset (if it has changed) should be provided.
- If this information is not available online, the authors are encouraged to reach out to the asset’s creators.

### 13. New assets

Question: Are new assets introduced in the paper well documented and is the documentation provided alongside the assets?

Answer: [Yes]

Justification: Original code and data are provided with documentation. Guidelines:

- The answer NA means that the paper does not release new assets.
- Researchers should communicate the details of the dataset/code/model as part of their submissions via structured templates. This includes details about training, license, limitations, etc.
- The paper should discuss whether and how consent was obtained from people whose asset is used.
- At submission time, remember to anonymize your assets (if applicable). You can either create an anonymized URL or include an anonymized zip file.

### 14. Crowdsourcing and research with human subjects

Question: For crowdsourcing experiments and research with human subjects, does the paper include the full text of instructions given to participants and screenshots, if applicable, as well as details about compensation (if any)?

Answer: [NA]

Justification: This research did not involve crowdsourcing nor human subjects. Guidelines:

- The answer NA means that the paper does not involve crowdsourcing nor research with human subjects.
- Including this information in the supplemental material is fine, but if the main contribution of the paper involves human subjects, then as much detail as possible should be included in the main paper.
- According to the NeurIPS Code of Ethics, workers involved in data collection, curation, or other labor should be paid at least the minimum wage in the country of the data collector.

### 15. Institutional review board (IRB) approvals or equivalent for research with human subjects

Question: Does the paper describe potential risks incurred by study participants, whether such risks were disclosed to the subjects, and whether Institutional Review Board (IRB) approvals (or an equivalent approval/review based on the requirements of your country or institution) were obtained?

Answer: [NA]

Justification: This research did not involve crowdsourcing nor human subjects. Guidelines:

- The answer NA means that the paper does not involve crowdsourcing nor research with human subjects.
- Depending on the country in which research is conducted, IRB approval (or equivalent) may be required for any human subjects research. If you obtained IRB approval, you should clearly state this in the paper.
- We recognize that the procedures for this may vary significantly between institutions and locations, and we expect authors to adhere to the NeurIPS Code of Ethics and the guidelines for their institution.
- For initial submissions, do not include any information that would break anonymity (if applicable), such as the institution conducting the review.

### 16. Declaration of LLM usage

Question: Does the paper describe the usage of LLMs if it is an important, original, or non-standard component of the core methods in this research? Note that if the LLM is used only for writing, editing, or formatting purposes and does not impact the core methodology, scientific rigorousness, or originality of the research, declaration is not required.

Answer: [NA]

Justification: LLMs were only used for minor editing and formatting. Guidelines:

- The answer NA means that the core method development in this research does not involve LLMs as any important, original, or non-standard components.
- Please refer to our LLM policy (https://neurips.cc/Conferences/2025/LLM) for what should or should not be described.

## Appendix

### A Derivation of the latent dynamics Eq. (5) from Eq. (4)

To go from Eq. (4) to Eq. (5), we only have to solve the integral

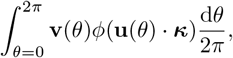

with **u**(*θ*) = (cos *θ*, sin *θ*)^T^ and **v**(*θ*) = *J*(cos(*θ* + Δ), sin(*θ* + Δ))^T^. (The symbol *·* above denotes the dot product.) Using the polar change of coordinate,

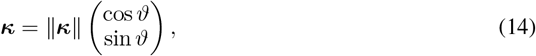

we have

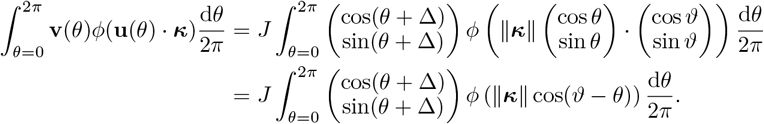

Recalling that *ϕ* is the Heaviside step function Θ, we have

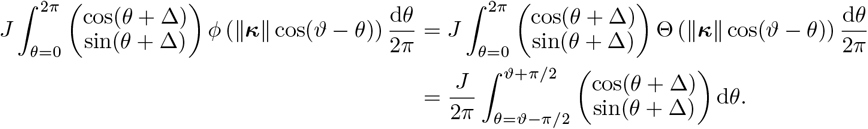

The integral above can be solved, which yields

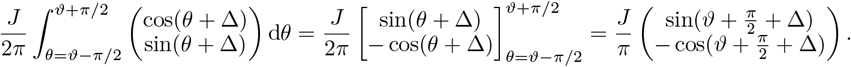

Using the sum and difference trigonometric identities,

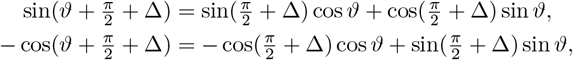

we can write

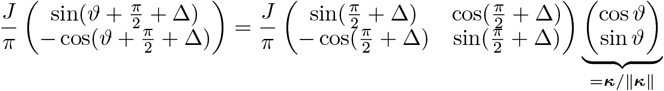

where we revert the polar change of coordinate, Eq. (14). Plugging in 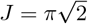 and Δ = *π/*4, we obtain

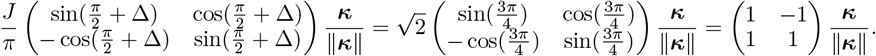

### B Derivation of pairwise correlations Eq. (6)

First, we compute the time-average and the standard deviation of the post-activation of neuron *i*,

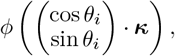

assuming that the latent variables ***κ*** rotate at constant speed on the unit circle (which is the asymptotic behavior of the dynamical system Eq. (5)). Using again the polar change of coordinate Eq. (14) and recalling that *ϕ* = Θ (the Heaviside step function), we have

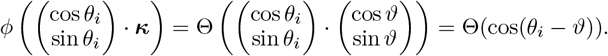

The time-average of the post-activation of the neuron *i* is

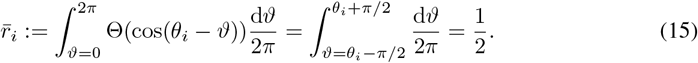

Similarly, the variance the post-activation of neuron *i* is

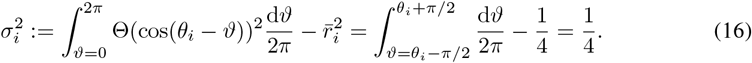

Hence, the correlation *C*_*ij*_ between any pair of neurons *i* and *j* is

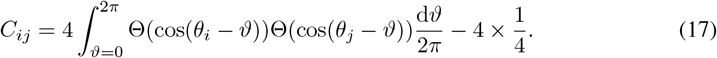

Developing the product in the integrand above, we see that it only remains to compute the integral

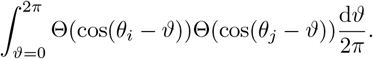

Using the change of variable 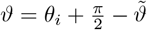, we have

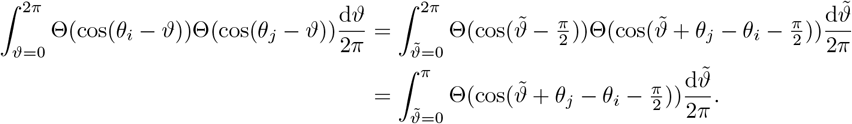

Without loss of generality, we assume that *θ*_*j*_ − *θ*_*i*_ ≥ 0 (otherwise, permute the indices *j* and *i*). We then distinguish two cases: If *θ*_*j*_ − *θ*_*i*_ ≤ *π*,

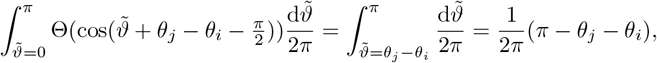

if *θ*_*j*_ − *θ*_*i*_ *> π*,

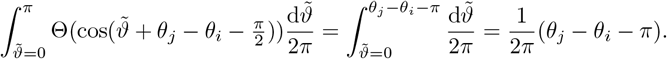

However, by the definition of the absolute angle difference |*θ*_*i*_− *θ*_*j*_ |:= cos^−1^(cos(*θ*_*i*_ − *θ*_*j*_)), we know that

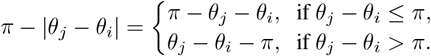

Hence,

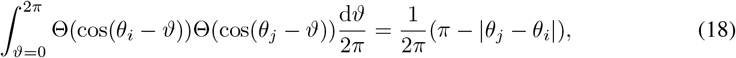

and we can verify that the equality above is invariant upon permutation of the indices *i* and *j*. Using Eqs. (15), (16), and (18) in Eq. (17), we obtain

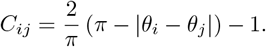

### C Simulation of an RNN with a finite number of neurons

To verify that the theory for infinite-size networks presented in Sec. 2 gives a good approximation of the activity of RNNs with a large but finite number of neurons, we simulated the dynamics of an RNN with *N* = 1000 neurons. The simulation shows that the latent variables ***κ*** are attracted to a limit cycle (Fig. 5A) and covariance of the network’s activity has a power-law eigenspectrum (Fig. 5A), as predicted by the theory for the infinite-size network.

**Figure 5.**
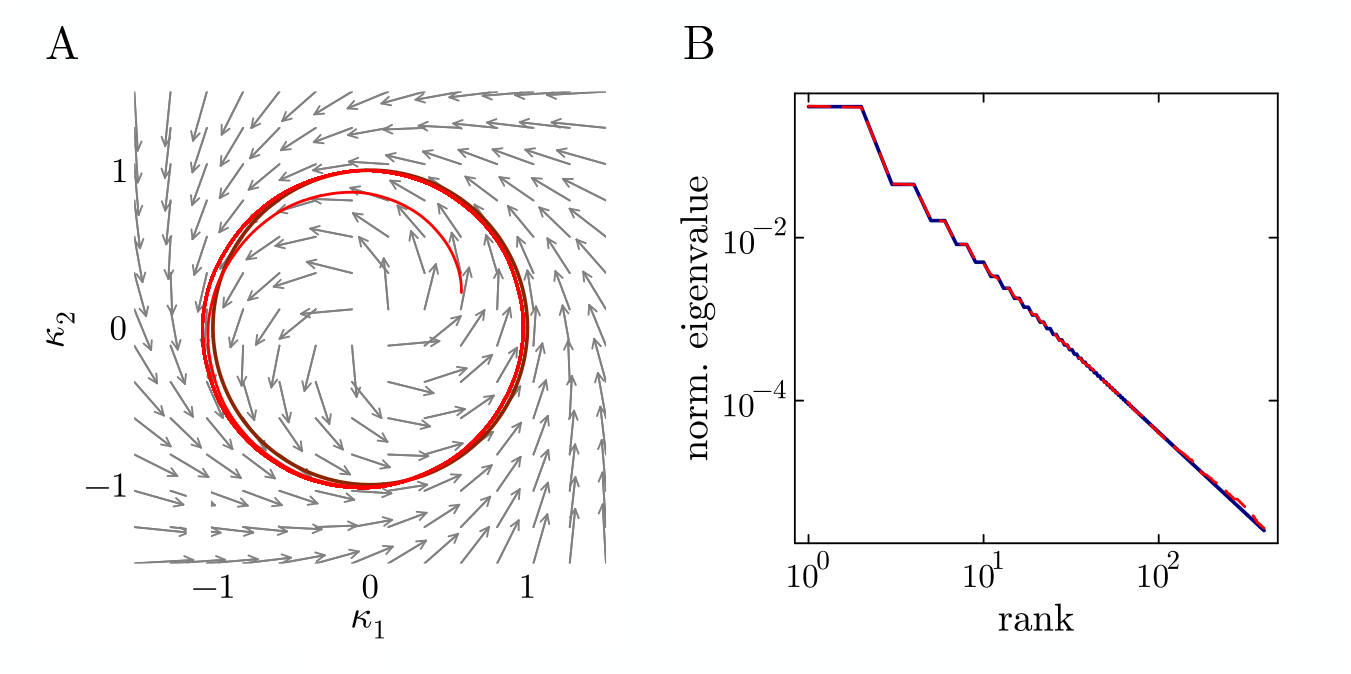
Simulation of an RNN with *N* = 1000 neurons. (**A**) Same as Fig. 1D, but with the addition of the trajectory (red curve) of the latent variables ***κ*** obtained from a simulation of the finite-size RNN. (**B**) Same as Fig. 1F, but with the addition of the PCA eigenspectrum (red dashed curve) obtained from the simulated activity of the finite-size RNN (only the first 400 eigenvalues are shown). The theory derived for infinite-size networks gives a good approximation of the activity of an RNN with *N* = 1000 neurons.

### D Solvable model with more general lateral connectivity

To illustrate that the analytical results obtained in Sec. 2 can be generalized to models with lateral connectivity that are more general than the rank-2, cosine connectivity prescribed by Eq. (2), we present here a solvable model where the weight matrix **W** is of rank 4 (inspired by model studied in [52]). Let us consider an RNN with *N* neurons and with a weight matrix given by

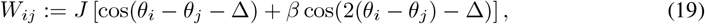

where the parameter *β* ≥ 0 fixes the importance of the higher frequency component, cos(2(*θ*_*i*_ − *θ*_*j*_) − Δ), of the connectivity. As in Sec. 2, the activation function *ϕ* is assumed to be the Heaviside step function Θ. Note that when *β* = 0, we recover the rank-2 RNN of Sec. 2. Following the same steps as in Sec. 2.1, and defining ***κ*** := **U**^†^**x**, the 4-dimensional vector of latent variables, we obtain that in the limit *N* → ∞, the dynamics of the latent variables ***κ*** follows

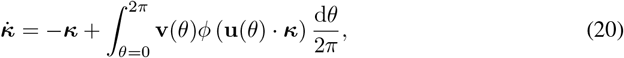

where

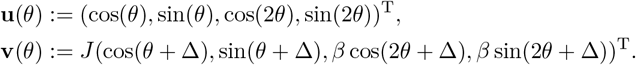

If we assume that at time *t* = 0, *κ*_3_(0) = *κ*_3_(0) = 0, i.e., the third and forth latent variables are null, we can use a polar change of coordinate similar to that used in Appendix A and write

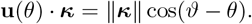

Then, recalling that *ϕ* = Θ, we can solve the integral in Eq. (20) and, choosing 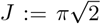 and Δ := *π/*4 as in Sec. 2, we find that

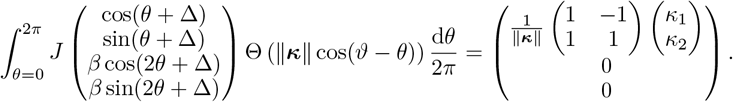

From the equality above, we deduce that *κ*_1_(*t*), *κ*_2_(*t*) follow the same dynamics as in the rank-2 network of Sec. 2, and *κ*_3_(*t*) = *κ*_4_(*t*) = 0, for all *t >* 0, i.e., the third and forth latent variables remain null at all times. Although we do not perform here a formal stability analysis of the limit cycle in the plane (*κ*_1_, *κ*_2_, 0, 0), we verified numerically that the latent variables of an RNN with *N* = 10^4^ neurons follow a limit cycle confined to the plane (*κ*_1_, *κ*_2_, 0, 0), when *β* is not too large (Fig. 6).

**Figure 6.**
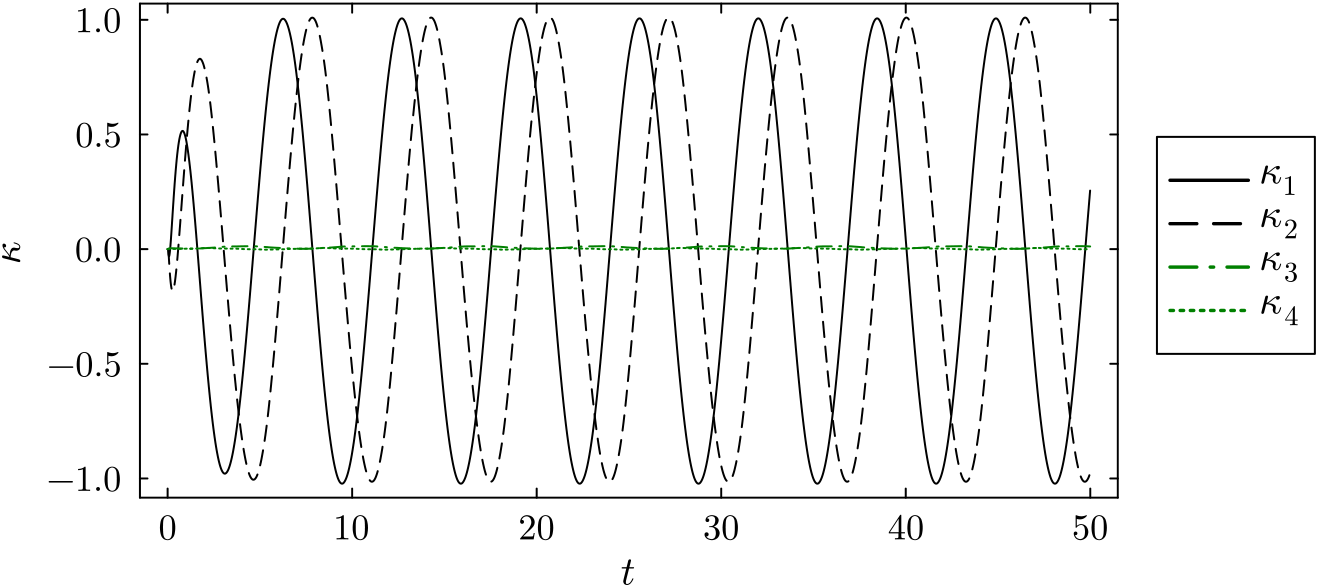
Simulation of the latent dynamics of an RNN with connectivity given by Eq. (19). In this simulation, *N* = 10^4^, *β* = 0.25, and all latent variables are initialized at 0 at time *t* = 0. The simulation shows that *κ*_1_ and *κ*_2_ converge to a limit cycle, whereas *κ*_3_ and *κ*_4_ remain close to **0**.

**Figure 7.**
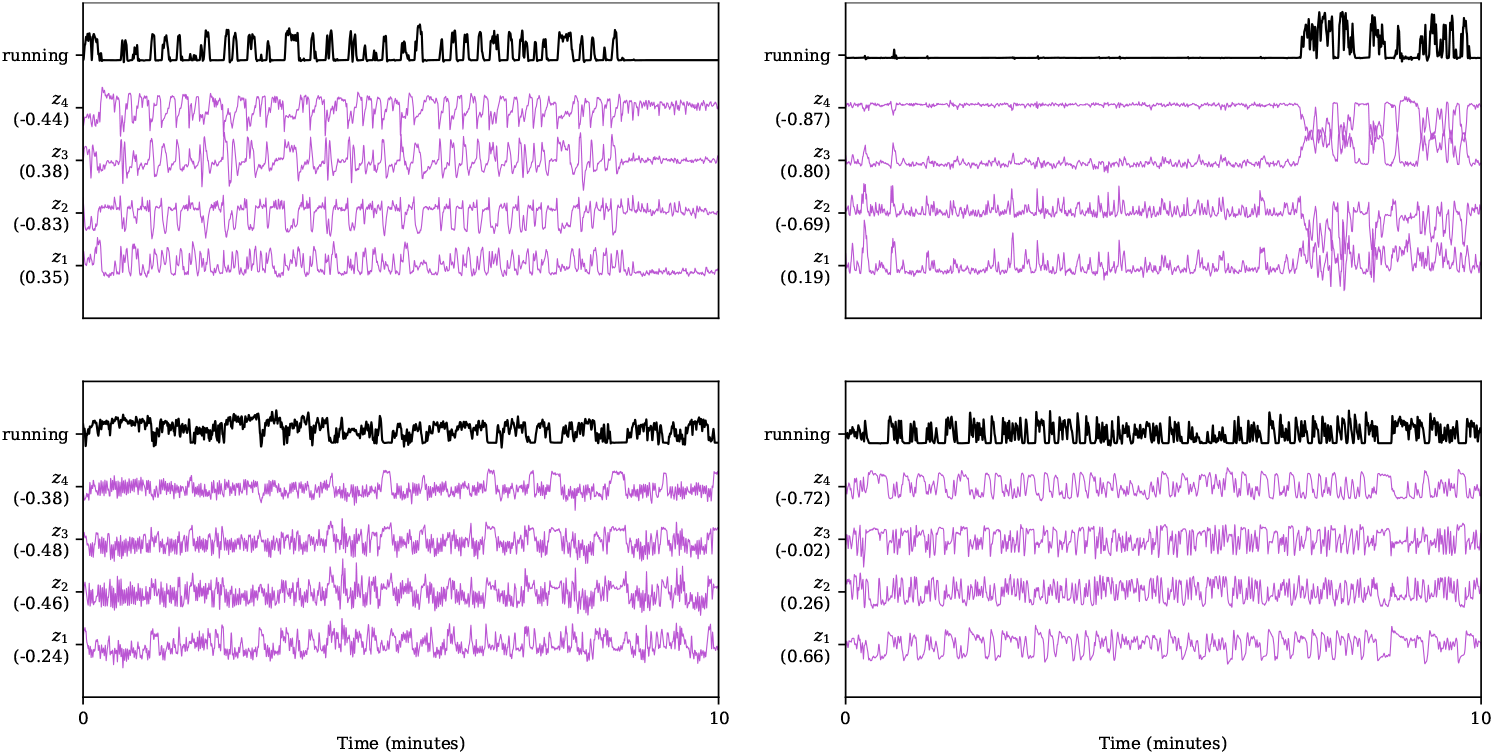
Extracted latents correlate with running speed. For four example sessions of spontaneous activity, the dynamics of the extracted latents *z*_*i*_ (purple) are correlated with the running speed (black). Correlation coefficient between each latent and running speed is reported in parentheses on the y-axis. All curves are smoothed with a 1s Gaussian kernel for visualization only.

Since the generalized model we have proposed can produce a limit cycle in the space or pre-activations that is identical to that of the rank-2 model of Sec. 2, we know that, when in this limit cycle, the post-activations of the neurons produce a power-law eigenspectrum identical to that of the rank-2 model. Hence, we have shown that the strong assumption of a rank-2, cosine weight matrix, Eq. (2), is not strictly necessary for an RNN to produce 2-dimensional latent dynamics and high-dimensional activity in the space of post-activations.

### E Random low-rank RNNs with tractable latent dynamics

The model analyzed in Sec. 2 is not the only model for which a simple expression for the latent dynamics can be obtained. To illustrate this fact, we re-use the general definition of RNN dynamics, Eq. (1), but consider here the class of low-rank RNNs with rank-*d* weights matrices **W** given by

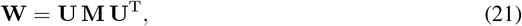

where **U** is an *N* × *d* random matrix with i.i.d. standard normal entries and **M** ∈ ℝ^*d*×*d*^ is an arbitrary “overlap” matrix [51]. As explained in [51], thanks to the fact that the “patterns” **U** are Gaussian, the dynamics of the *d*-dimensional latent variable vector ***κ*** := **U**^†^**x** can be described, in the large-network limit *N* → ∞, by the dynamical system

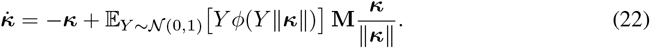

In the present work, we go one step further and ask whether the expectation 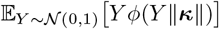 can be solved, in order to yield simpler expressions for the latent dynamics. The following proposition shows that the answer is positive when the activation function *ϕ* is either (i) a step function, (ii) a ReLU function, or (iii) a Gaussian cumulative distribution function

#### Proposition 1.

*(i) For any bias b* ∈ ℝ, *if ϕ is the step function ϕ* = Θ(*x* + *b*), *Eq*. (22) *reduces to*

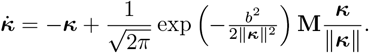

*(ii) For any bias b* ∈ ℝ, *if ϕ is the ReLU function ϕ* = max(0, *x* + *b*), *Eq*. (22) *reduces to*

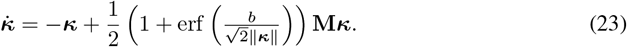

*(iii) For any mea n µ* ∈ ℝ *and variance σ*^2^ ≥ 0, *if ϕ is the Gaussian c*.*d*.*f*. 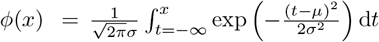, *Eq*. (22) *reduces to*

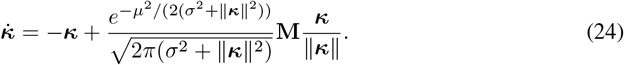

The proposition shows the nontrivial effect of the activation function has on the latent dynamics. For example, let us take again *d* = 2. When *ϕ* is the step function with bias *b* = 0, and if 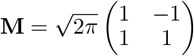, we recover the latent dynamics Eq. (5), which generates a stable limit cycle.

In contrast, when *ϕ* is the ReLU function with bias *b* = 0, Eq. (23) becomes the linear system 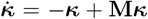, which can generate rotational dynamics but cannot generate a stable limit cycle. Note that Eq. (24) generalizes a previous result obtained for the erf activation function, used as an approximation of tanh [74].

*Proof*. The prove these statements, it suffices to solve the integral,

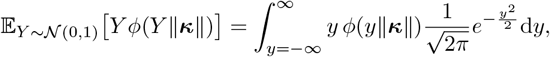

for the three different functions *ϕ*.

For *(i)*,

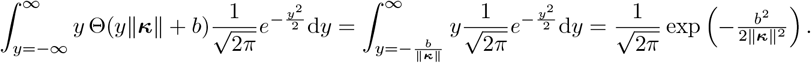

For *(ii)*,

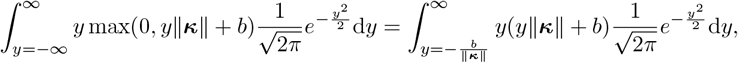

and we integrate by parts,

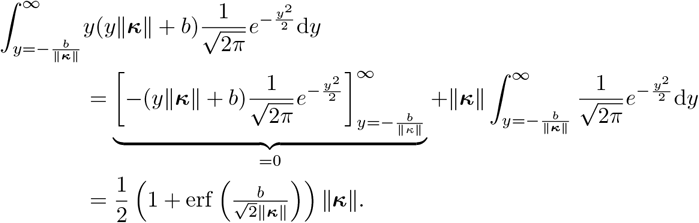

For *(iii)*, we first integrate by parts,

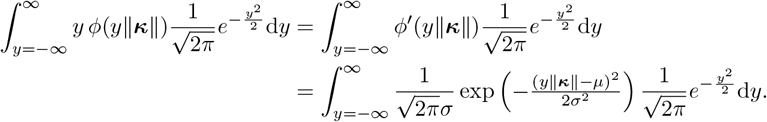

Then, we complete the square,

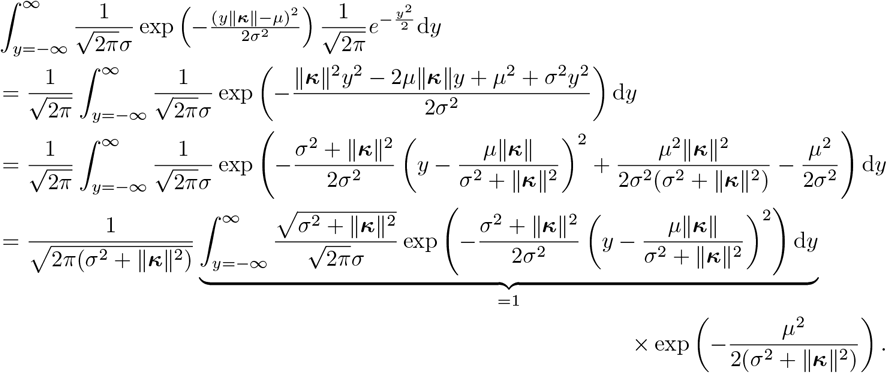

### F From the eigenvalues of the activity matrix A^(*N,T*)^ to the eigenvalues of the random feature kernel operator Eq. (10)

First, we recall that the eigenvalues 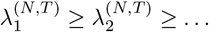 of the “covariance matrix”,

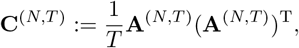

are related to the eigenvalues 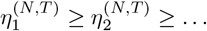 of the “Gram matrix”,

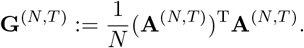

Indeed, using the singular value decomposition of **A**^(*N,T*)^, one can easily verify that

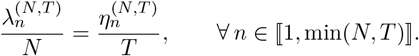

By the law of large numbers, in the limit *N* → ∞, the entries of the Gram matrix **G**^(*N,T*)^ converge to an expectation corresponding to the random feature kernel Eq. (11): For any 1 ≤ *s, t* ≤ *T*,

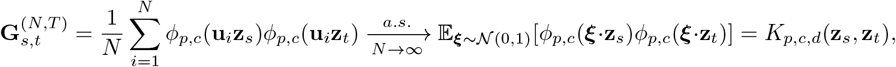

where **u**_*i*_ above denotes the *i*-th row of the *N* × *d* random matrix **U**. Hence, as *N*→ ∞, the eigenvalues 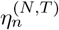 of the Gram matrix **G**^(*N,T*)^ converge to the eigenvalues 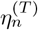 of the kernel matrix

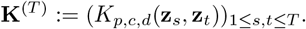

Furthermore, we have that

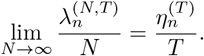

Finally, by a theorem from Koltchinskii and Giné [54, Theorem 3.1], the scaled eigenvalues 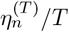 of the of the kernel matrix **K**^(*T*)^ converge (in a ℓ_2_ sense) to the eigenvalues *λ*_*n*_ of the integral operator Eq. (10). In conclusion, we have that, for any given *n*,

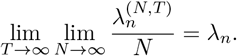

### G Gaussian processes from RNNs

As in Appendix B, we start from an infinite-size network where the latent variables ***κ*** rotate at constant speed on the unit circle. Using again the change of coordinate Eq. (14), the post-activation (firing rate) of a neuron with location *θ*_*i*_ is

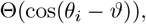

where *ϑ* is the angle of the latent variable ***κ***. Also, from Eq. (5), one can easily deduce that the norm of the velocity vector is always 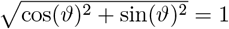, which implies that the cycle time period of the system is 2*π*.

Taking *M* independently and uniformly sampled neurons on the circle, we consider the random projection readout

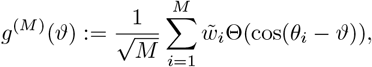

where the readout weights 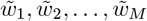 are *i*.*i*.*d*. standard normal variables that are also independent from the random angles *θ*_*i*_. Since the readout *g*^(*M*)^ is a weighted sum of 2*π*-periodic functions, *g*^(*M*)^ is itself a 2*π*-periodic function.

As the number of randomly sampled neurons *M* tends to infinity, the random readout *g*^(*M*)^ converges in law to a Gaussian process on the circle 𝕊^1^. To see this, we first compute the mean and covariance of the 2*π*-periodic function *g*^(*M*)^, as *M* → ∞. For the mean, we have

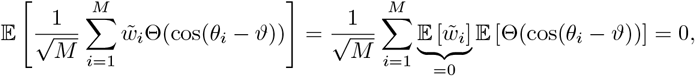

for any *M* . For the covariance, we have, we have, for any pair of angles *ϑ* and *ϑ*^*′*^,

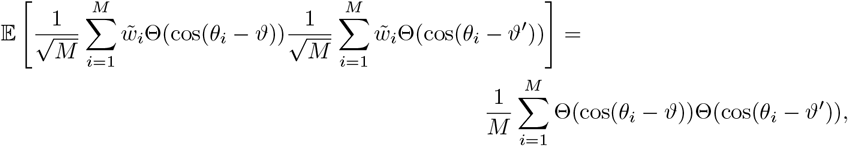

using that 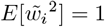 and 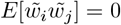 when *i* ≠ *j*. By the law of large numbers,

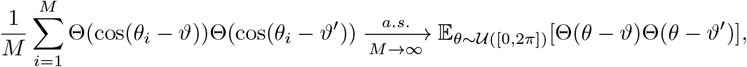

and using the same line of arguments we used to derive Eq. (18) in Appendix B, we obtain

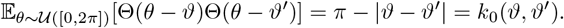

Then, by the multivariate central limit theorem, we have that, for any finite sequence of angles *ϑ*_1_, *ϑ*_2_, …, *ϑ*_*m*_,

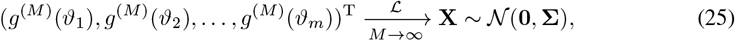

where

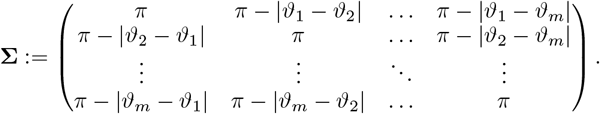

By the Kolmogorov extension theorem, there exists a Gaussian process on 𝕊^1^, *g* ∼𝒢𝒫 (**0**, *k*_0_), such that **X** can be replaced by (*g*(*ϑ*_1_), *g*(*ϑ*_2_), …, *g*(*ϑ*_*m*_))^T^ in Eq. (25), for any finite sequence of angles *θ*_1_, *θ*_2_, …, *θ*_*m*_. Hence, we have shown that the random projection readout *g*^(*M*)^ defines a random 2*π*-periodic function that has the same law as a Gaussian process as *M* → ∞.

In summary, the above arguments show that our solvable RNN model can be used to draw samples from a periodic Gaussian process. In other words, it can represent a Gaussian process prior over periodic functions. This indicates that the Gaussian process framework originally developed for infinite-width feedforward neural networks [87, 88] also applies, in certain cases, to large networks with recurrent dynamics.

### H *In vivo* experimental methods

#### Neuronal Recordings

Transgenic mice expressing the GCamP6s calcium indicator [94] in all excitatory cortical neurons (CamKII x Ai162) were implanted with a 4 mm imaging window over visual cortex. Mice were headfixed under a custom Light Beads Microscope [65], modified with an optical stabilization module. A volumetric field of view of 2 × 2 × 0.3 mm^3^ over primary and higher visual areas was imaged with 2-3 µm lateral and 20 µm axial voxel spacing, with volume rates between 4.1-4.5 Hz. During recordings, mice were not anesthetized and were free to run on a low-resistance treadmill.

#### Visual Stimulation

Visual stimuli were delivered via three screens that covered approximately 270^*?*^ horizontally and 70^*?*^ vertically of the visual field of the mouse. During drifting grating experiments, full-field drifting gratings with a temporal frequency of 2.0 Hz and spatial frequency of 0.04 cycles/degree were presented for 2 s, with an inter-stimulus interval (ISI) drawn from a uniform distribution between 2-3 s. A sequence of 720 stimuli was presented with random directions drawn from a uniform distribution of integer angles ranging from 0^*?*^ to 359^*?*^. The same set of 720 stimuli was repeated in a new random order in the second half of the experiment. For natural image experiments, 1866 unique stimuli were presented, each composed of a mosaic of three distinct natural images. The images were chosen from the same set presented in [39]. Each stimulus was presented for 0.8 s with a random ISI between 0.9-1.3 s. Stimuli were presented in two shuffled sequences such that each stimulus was presented twice. For sessions of spontaneous activity, no stimuli were displayed and the screens were grey. Mice were awake, free to run (as in the other sessions) for 10 to 15 minutes of imaging.

#### Data preprocessing

To detect cells and extract neuronal activity from volumetric fluorescence movies, we used Suite3D [66], an analysis pipeline for motion-correction, cell detection and signal extraction from volumetric two-photon data. After semi-automated curation, we analyzed activity from 19,223 ± 2,948 (mean ± std) neurons per experiment, across 10 experiments (4 spontaneous activity, 3 natural images, 3 drifting gratings, across three mice). The extracted cell fluorescence was deconvolved after neuropil substraction. Deconvolved traces were re-sampled at 5 Hz and aligned to stimulus onset events. For stimulus response sessions, we projected out the activity along the top 30 principal components (PCs) of spontaneous activity (computed on a spontaneous session recorded consecutively from the same mouse), similar to what was done in [39]. To compute stimulus responses, neuronal activity within a response window was averaged (0.5 s for images, 2.0 s for gratings) to produce a single value per neuron per stimulus presentation. Deconvolved spontaneous activity was resampled using linear interpolation at 5 Hz.

The recordings covered part of primary visual cortex, higher visual areas AM, PM, LM, and RL, with occasional inclusion of some somatosensory areas. We confirmed the recorded area via retinotopic mapping using sparse noise stimuli. To determine stimulus responsiveness, we used the signal-related variance metric to determine what fraction of a neuron’s variance could be explained by stimuli (described in Stringer et al. Nature 2019). In a typical recording with 22,000 neurons, the top 1000 cells had 49% stimulus-related variance, the next 1000 had 22%, 14%, 10% and so on, where the 10,000th cell had 3%. The responsive neurons were distributed across primary and higher visual cortices, and the selection of source and target neurons was agnostic to location.

### I Neural Cross-Encoder

#### I.1 Datasets

For all three data conditions (gratings, natural images and spontaneous activity), we compared the performance of the (nonlinear) NCE model to (linear) Reduced Rank Regression on the same datasets. For results in Fig. 4 we used datasets with 500 source neurons and 1,000 target neurons. For stimulus-driven conditions (gratings and images), the source neurons were chosen to be the 500 neurons with the most reliable stimulus-evoked responses, quantified by the fraction of stimulus-related variance defined in Ref. [39]. 1,000 target neurons were chosen from the 2,000 non-source neurons with the highest stimulus-related variance. For the spontaneous condition, the target and source neurons were selected at random from the full population. We repeated the model fit for each condition and session five times with random initializations and neuron selections. In all cases, source and target neuron sets were non-overlapping.

To reduce the impact of non-stimulus related activity in stimulus-driven conditions, we used the fact that each stimulus was presented multiple times, and trained the model to predict the activity of the target neurons on one repeat of a stimulus from the activity of the source neurons on a different repeat, using all possible permutations. For example, if the same stimulus was shown at times *t*_1_, *t*_2_, the network was trained to minimize the sum of the squared Euclidean distances 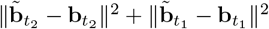, where 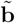 is the recorded activity and **b** is the NCE reconstruction.

For the spontaneous condition, to capture the temporal structure in the latent dynamics, we used multiple timepoints of the source neurons to estimate the latent variable at a single timepoint. To predict **z**_*t*_, we took a window of size *L* timepoints and concatenated the source activity within this time window, increasing the input dimensionality of the encoder: 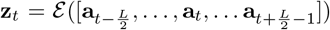. The number and size of the hidden layers were unchanged. For the experiments presented, the time window was 1.2 s (*L* = 6 timepoints).

In all three conditions, the paired source-target datasets were split into train-validation-test sets with a 50%-20%-30% split. For stimulus-driven datasets, the split was done using stimulus identity to ensure that all permuted pairs of repeats of a single stimulus were in the same set after splitting. For spontaneous datasets, instead of assigning each timepoint to one of three sets independently, we separated the time series into 10 s chunks with 2 s buffers between each chunk, and assigned each chunk to one of three sets. This procedure prevents the training set from contaminating the test set through the slow temporal autocorrelation of calcium signals.

#### I.2 Model Fitting

Linear models were fit on the training set using the closed-form solution of Reduced Rank Regression with a ridge penalty. The ridge penalty (which varied between 10^−1^-10^6^) giving the best performance in the validation set was selected.

For the NCE, the encoder is a 3-layer feedforward network with ReLU activation functions. The hidden layers contain (500, 250, 100) units. The parameters of the encoder *E* and the parameters of the decoder, i.e., the readout weights and biases **U** ∈ ℝ^*B*×*d*^ and **c** ∈ ℝ^*B*^, the activation parameter *p*, and the baseline firing rates **r**_0_ ∈ ℝ^*B*^, were learned by minimizing the mean-squared error (MSE) loss 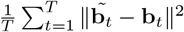, where **b**_*t*_ = *ϕ*_*p*,**c**_(**U**ℰ (**a**_*t*_)) + **r** (**b**_*t*_ is the predicted activity of the target neurons, **a**_*t*_ is the recorded activity of the source neurons, and ℰ represents the encoder). NCE models were trained in two phases: pretraining and fine-tuning. In the pretraining phase, we used a form of knowledge distillation [95]. Instead of predicting the activity of target neurons directly, the NCE was trained on the predicted activity of target neurons from the best-fitting linear model, one that potentially had a higher pre-activation dimension, *d*, than the NCE. In the fine-tuning phase, the NCE was trained to predict the true neuronal activity.

In both phases of training, we augmented the training set by adding noise to the activity of source neurons. First, we added Poisson-like shot noise present in two-photon recordings by generating zero-mean Gaussian noise with a variance equal to the activity of each neuron at a given timepoint, scaled by a factor of 0.05. Next, we added independent Gaussian noise with a mean of 0 and variance of 0.05. Finally, we incorporated multiplicative noise, multiplying the activity of each neuron at each timepoint with an independent Gaussian random variable with a mean of 1 and variance of 0.05. The original source dataset was concatenated with three noise-augmented datasets to produce the final training set.

In each phase, parameters were optimized through stochastic gradient descent using Adam [96] and a batch size of 2,048, with a learning rate of 10^−3^, momentum parameters *β*_1_ = 0.85, *β*_2_ = 0.95, and an *L*_2_ penalty of 10^−5^. Model parameters were randomly initialized with Kaiming initialization [97]. Within each phase, we iterated between optimizing the readout parameters, encoder parameters, and all parameters. At each iteration, the model with the best validation performance, evaluated using the coefficient of determination, was selected. Once models were fitted, we report the fraction of explained variance on the test set. To normalize this value to only account for the “explainable” variance, we computed the coefficient of determination of each model, and normalized by the maximum of the best model across all pre-activation dimensionalities for the given set of neurons.

#### I.3 Hyperparameter Selection

The architecture of the encoder, the numbers of source and target neurons, as well as the training parameters were selected to balance computational efficiency with performance. The encoder is a feedforward network with parametrized by the number of layers (depth) and the number of units in the first layer (width), with the width of each successive layer reduced by a factor of two. We conducted grid searches on example sessions across a range of values of *d* and found that the resulting performance is only mildly sensitive to the encoder of the architecture (see Tables 1,2, 3, 4 for representative examples).

**Table 1:** Normalized *R*^2^ for an example grating session with *d* = 4.

| Layer size \ # Layers | 3 | 2 | 1 |
| --- | --- | --- | --- |
| 1000 | 0.97 | 0.97 | 0.96 |
| 500 | 0.97 | 0.97 | 0.94 |
| 250 | 0.95 | 0.96 | 0.94 |

**Table 2:** Normalized *R*^2^ for an example natural image session with *d* = 8.

| Layer size \ # Layers | 3 | 2 | 1 |
| --- | --- | --- | --- |
| 1000 | 0.54 | 0.55 | 0.52 |
| 500 | 0.55 | 0.56 | 0.53 |
| 250 | 0.56 | 0.54 | 0.51 |

**Table 3:** Normalized *R*^2^ for an example natural image session with *d* = 32.

| Layer size \ # Layers | 3 | 2 | 1 |
| --- | --- | --- | --- |
| 1000 | 0.91 | 0.89 | 0.89 |
| 500 | 0.91 | 0.92 | 0.89 |
| 250 | 0.92 | 0.91 | 0.91 |

**Table 4:** Normalized *R*^2^ for an example natural image session with *d* = 64.

| Layer size \ # Layers | 3 | 2 | 1 |
| --- | --- | --- | --- |
| 1000 | 0.97 | 0.98 | 0.96 |
| 500 | 0.95 | 0.92 | 0.98 |
| 250 | 0.98 | 0.99 | 1.00 |

Due to computational constraints, we subset the full population of neurons into 1000 source and 500 target neurons. Varying these numbers up to 2000 neurons does not substantially affect the results shown in Figure 4. Tables 5, 6 reports the fraction of explainable variance explained (normalized *R*^2^) for a representative example session of grating responses, fit with varying subset sizes. Increasing the number of cells beyond 2000 somewhat reduces performance and we suggest two potential explanations. First, due to animal health concerns our experiments are of limited duration, hence a typical session produces only 1000 training examples. This may not be enough to train a relatively large encoder network with >1000 input neurons. Second, only a fraction of neurons recorded have reliable stimulus responses, and including more cells introduces a large amount of non-stimulus-related variance.

**Table 5:** Normalized *R*^2^ with an NCE model for an example grating session with *d* = 4.

| # Target \ # Source | 100 | 500 | 1000 | 2000 |
| --- | --- | --- | --- | --- |
| 100 | 0.98 | 0.96 | 0.96 | 0.97 |
| 500 | 0.98 | 0.97 | 0.94 | 0.93 |
| 1000 | 0.98 | 0.97 | 0.94 | 0.91 |
| 2000 | 0.99 | 0.97 | 0.97 | 0.92 |

**Table 6:**
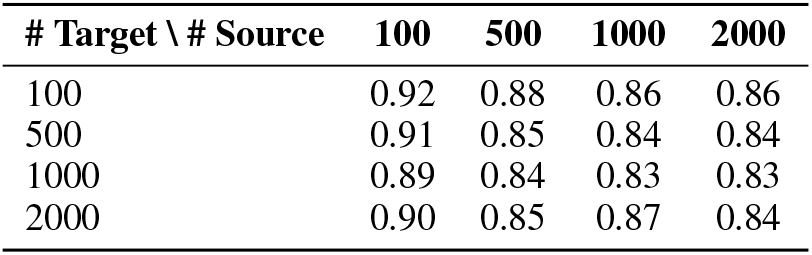
Normalized *R*^2^ with an RRR model for an example grating session with *d* = 4.

For the remaining hyperparameters, we explored the hyperparameter space with the Optuna optimizer (Akiba et al. KDD 2019). We found that the performance of the NCE is not sensitive to modest variations in learning rate, batch size, and momentum.

#### I.4 Latent dynamics of spontaneous activity

To determine whether the extracted latent variables related to the behavioral state of the animal, we compared them to the running speed of the animal. For each session of spontaneous activity, we trained an NCE model with a pre-activation dimension of 4, as described above. After training, we took the activity of the source neurons across the entire session (removing the train/validation/test splits), and computed the inferred latents.

The running speed of the mouse is highly correlated with inferred latent variables. We computed the correlation coefficient between each latent and the running speed, and found high positive or negative correlations for most variables. This supports the hypothesis that the latents inferred by the NCE are related to the behavioral state of the animal.

#### I.5 Compute Resources

Volumetric calcium movies were processed using a custom workstation running Ubuntu 20.04.6 LTS with an Intel(R) Xeon(R) w9-3475X CPU (36 cores, 2.20 GHz), NVIDIA RTX A4500 GPU (20GB VRAM), and 512 GB DDR5 RAM. Preprocessing with Suite3D took <20 hours per session.

All NCE experiments were performed on a workstation running Windows 10 Pro, with an Intel (R) Core(TM) i7-11700k CPU (8 cores, 3.6 GHz), NVIDIA RTX 3060 GPU (12GB VRAM), and 128 GB DDR4 RAM. Total runtime of all NCE experiments was <96 hours. All preliminary experiments that are not reported in the paper were conducted on this workstation, with similar runtimes.

#### I.6 Software and Licenses

This work makes use of several publicly available software resources including: Python (PSF License), Matplotlib (PSF License), NumPy (BSD License), SciPy (BSD License), CuPy (MIT License), PyTorch (BSD-3 License), Suite3D (AGPL-3 License).

#### I.7 Code and Data Availability

All data and code used to train the NCE and generate the results, as well as code used for simulations in Fig. 2, are shared via Figshare: https://figshare.com/s/aac82eac1829b7ec406e and Github: https://github.com/alihaydaroglu/NCE.

## Footnotes

2 cos(*α* − *β*) = cos(*α*) cos(*β*) + sin(*α*) sin(*β*)

3 The *k*-th weak derivative of a function *f* : ℝ → ℝ is the defined as the function 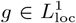 that satisfies 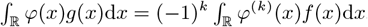, for all 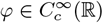.

